# Age-associated sleep-wake patterns are altered with Prdm13 signaling in the dorsomedial hypothalamus and dietary restriction in mice

**DOI:** 10.1101/2022.09.26.509442

**Authors:** Shogo Tsuji, Cynthia S Brace, Ruiqing Yao, Yoshitaka Tanie, Hirobumi Tada, Nicholas Rensing, Seiya Mizuno, Julio Almunia, Yingyi Kong, Kazuhiro Nakamura, Noboru Ogiso, Shinya Toyokuni, Satoru Takahashi, Michael Wong, Shin-ichiro Imai, Akiko Satoh

**Author notes:** Corresponding author: Akiko Satoh, Ph.D., Associate Professor, Department of Integrative Physiology, Geroscience Research Center, National Center for Geriatrics and Gerontology, 7-430 Morioka-cho, Obu, Aichi, 474-8511, Japan. equally contributing authors (Shogo Tsuji, Cynthia S Brace).

## Abstract

Old animals display significant alterations in sleep-wake patterns such as increases in sleep fragmentation and sleep propensity. Here we demonstrated that dorsomedial hypothalamus-specific *PR-domain containing protein 13*-knockout (DMH-*Prdm13*-KO) mice recapitulated age-associated sleep alterations such as sleep fragmentation and increased sleep attempts during sleep deprivation (SD). These phenotypes were further exacerbated during aging, with increased adiposity and decreased physical activity, resulting in shortened lifespan. Dietary restriction (DR), a well-known anti-aging intervention in diverse organisms, ameliorated age-associated sleep alterations, whereas these effects of DR were abrogated in DMH-*Prdm13*-KO mice. Moreover, overexpression of *Prdm13* in the DMH ameliorated sleep fragmentation and excessive sleepiness during SD in old mice. Therefore, maintaining Prdm13 signaling in the DMH might play an important role to control sleep-wake patterns during aging.

---

The elderly commonly experiences changes in their sleep habits and sleep disruptions that causes problems, including waking often during the night, needing daytime naps, and having trouble falling asleep. The National Institute on Aging conducted a multicentered study called “Established Populations for Epidemiologic Studies of the Elderly (EPESE)” with more than 9,000 participants aged 65 years and older^1^. Interestingly, people who reported excessive sleepiness during the afternoon or evening had a slight, but statistically significant increase in the odds for 3-year mortality. Even in mice, recent studies have demonstrated that old C57BL/6J mice exhibit reduced amount of wakefulness and increased amount of non-rapid eye movement (NREM) sleep^2-4^. In both humans and mice, old individuals display sleep fragmentation, characterized by shorter episode durations of wakefulness, NREM and REM sleep, compared with young individuals^2,3,5-8^. Indeed, chronic sleep fragmentation is known to be associated with derailments in physiological functions, including low physical activity, increased adiposity and metabolic dysfunction^9,10^. Thus, it is conceivable that the dysregulation of sleep-wake patterns has a mechanistic connection to age-associated physiological decline. However, such a mechanistic connection has remained elusive, and it is unclear whether any effective intervention could improve age-associated sleep dysfunction.

The hypothalamus plays a critical role in the regulation of sleep-wake patterns^11^ and aging and longevity in mammals^12-14^. In our previous study, we have demonstrated that the mammalian NAD^+^-dependent protein deacetylase Sirt1 in the dorsomedial and lateral hypothalami (DMH and LH, respectively) delays aging, with significant enhancement of physical activity, oxygen consumption, body temperature and delta power, which is an indicator for depth of sleep, and extends lifespan in mice^12-14^. Furthermore, knockdown of *Sirt1* in the DMH and LH causes low delta power, and another mouse model with high hypothalamic Sirt1 activity displays reduced sleep fragmentation with advanced age and lifespan extension^15^. These findings suggested a possibility that a specific subpopulation of Sirt1-expressing neurons in the DMH and/or LH controls sleep-wake patterns during the process of aging. We previously conducted a comprehensive transcriptome analysis to identify DMH-enriched genes and identified *PR-domain containing factor 13 (Prdm13)*. Importantly, *Prdm13* is regulated by Sirt1 signaling, and DMH-specific *Prdm13* knockdown mice show low delta power^16^. Therefore, we hypothesized that Prdm13 signaling in the DMH, where Sirt1 signaling is involved in aging and longevity control, is causally involved in age-associated sleep alterations.

In the present study, to address this hypothesis, we generated DMH-specific *Prdm13*-knockout (DMH-*Prdm13*-KO) mice and found that DMH-*Prdm13*-KO mice display sleep fragmentation and excessive sleepiness during sleep deprivation (SD), which are common phenomena in aged C57BL/6J mice. Aging DMH-*Prdm13*-KO mice displayed further exaggerated sleep alterations, increased adiposity, decreased physical activity and shortened lifespan. We also found that dietary restriction (DR), a well-known anti-aging intervention in diverse organisms^17^, ameliorates age-associated derailment of sleep-wake patterns. These effects of DR were abrogated in DMH-*Prdm13*-KO mice. Moreover, overexpression of *Prdm13* in the DMH ameliorated sleep fragmentation and excessive sleepiness during SD in old mice. Thus, our results suggest that Prdm13 is involved in the regulation of sleep-wake patterns by DR, and that maintaining the function of Prdm13 signaling promotes youthful sleep-wake patterns during the process of aging.

## Results

### Old mice showed increases in sleep fragmentation and sleep propensity compared to young mice

Sleep fragmentation is one of the most common clinical characteristics in old individuals in both humans and mice^2,3,5-8^. To confirm age-associated sleep fragmentation, we conducted electroencephalogram (EEG) and electromyogram (EMG) recordings in young and old mice at 4 and 20 months of age, respectively. Old mice displayed greater sleep fragmentation compared to young mice during the light period (rest period), and predominantly during the dark period (active period) (**Fig. 1a,b**). During the dark period, the number of wakefulness, NREM and REM sleep episodes in old mice were significantly higher than young mice (**Fig. 1a**), whereas the duration of wakefulness and REM sleep episodes in old mice were shorter than young mice (**Fig. 1b**). The duration of REM sleep episodes in old mice was also shorter during the light period (**Fig. 1b**). The number of wakefulness episodes in old mice was significantly higher than young mice during the light period at ZT6-8, whereas the number of NREM sleep episodes in old mice was tended to be higher (**Fig. 1a**). In addition, in a 24-hour period, old mice spent less time awake and more time in NREM sleep (**Fig.1c, Supplementary Fig. 1a**). As most mouse studies reported^2-4,7^, the total amount of wakefulness in old mice was significantly lower than young controls during the dark period, whereas the total amount of NREM sleep was higher (**Fig. 1c**). Similar differences were also observed during the light period (**Fig.1c**). Together, our data confirm that old mice display greater sleep fragmentation and spend more time asleep compared to young mice.

**Fig. 1:**
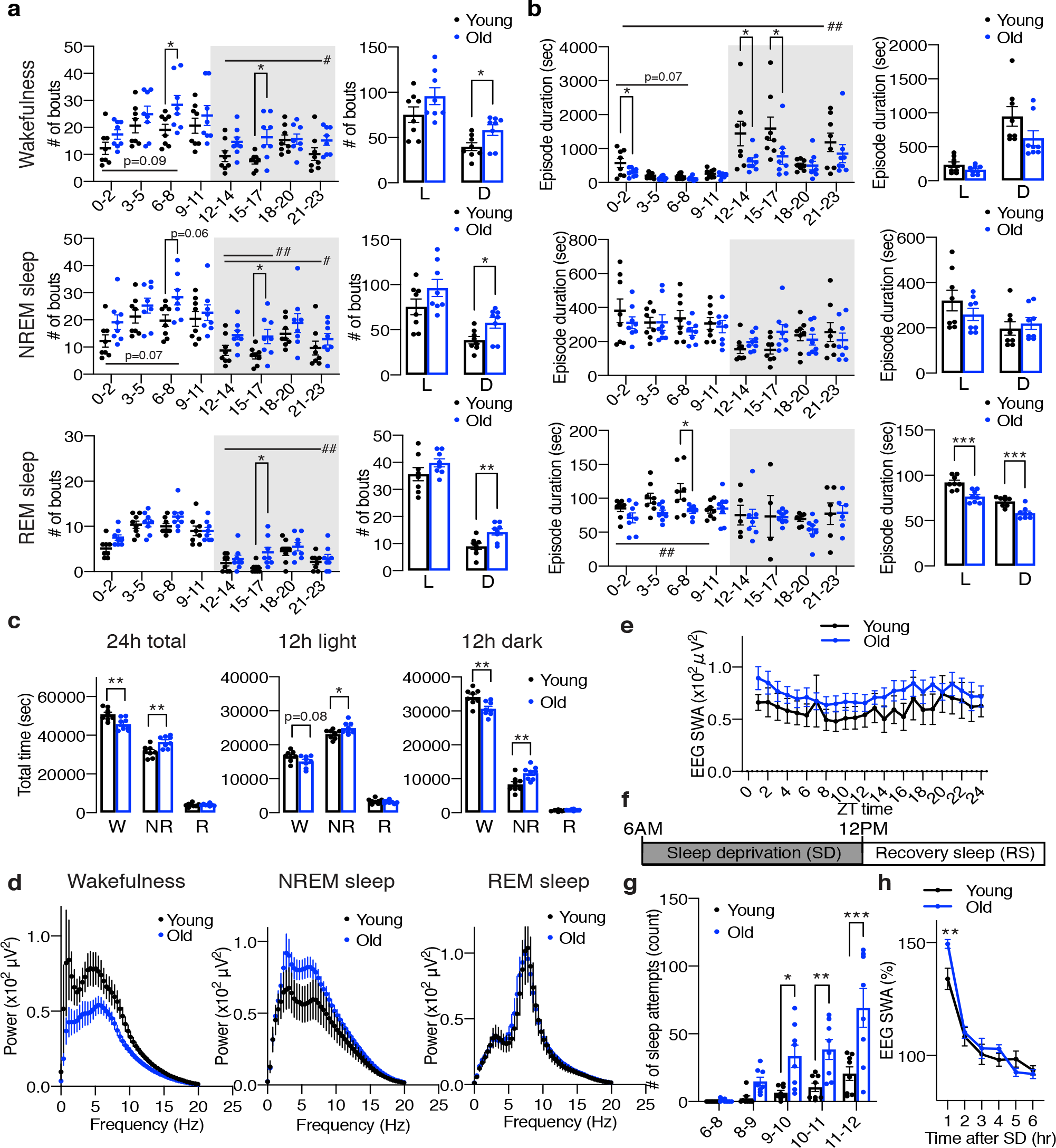
Old C57BL/6J mice display increases in sleep fragmentation and sleep propensity, low NREM EEG spectra, and excessive sleepiness during SD. **a**,**b**. Numbers of episodes (**a**) and duration (**b**) of wakefulness (top), NREM sleep (middle) and REM sleep (bottom) every 3 hours through a day (left) and during the light (L) and dark (D) periods (right) in young and old mice (n=8). Shading indicates dark period. Values are shown as means ± S.E., #p<0.05 and ##p<0.01 by repeated measures ANOVA, listed p-values, *p<0.05, **p<0.01 and ***p<0.001 by repeated measures ANOVA with Bonferroni’s post hoc test (left) or unpaired t-test (right). **c**, Total amount of wakefulness, NREM sleep and REM sleep during a 24-hour period (24h total), 12-hour light period (12h light) or 12-hour dark period (12h dark) (n=8). Values are shown as means ± S.E., listed p-values, *p<0.05 and **p<0.01 by unpaired t-test. **d**, EEG spectra of wakefulness (left), NREM sleep (middle) and REM sleep (right) during the light period (n=7-8). Values are shown as means ± S.E. **e**, SWA in the range of frequencies between 0.5 to 4 Hz during NREM sleep for a 24-hour period (n=7-8). Values are shown as means ± S.E. **f**, Schematic of SD. Sleep was deprived for six hours, between 6am and 12pm, followed by a period of RS. **g**, Number of sleep attempts during SD from 6am to 8am (6-8), 8am to 9am (8-9), 9am to 10am (9-10), 10am to 11am (10-11) and 11am to 12pm (11-12) in young and old mice (n=8). Values are shown as means ± S.E., *p<0.05, **p<0.01 and ***p<0.001 by repeated measures ANOVA with Bonferroni’s post hoc test. **h**, SWA after SD in young and old mice (n=8). Each value is relative to the average of the 24-hour baseline day. Values are shown as means ± S.E., **p<0.01 by repeated measures ANOVA with Bonferroni’s post hoc test.

To examine the profile of EEG spectra during each state in young and old mice, we used fast Fourier transform (FFT) of EEG recordings. During wakefulness, the power of EEG in the frequency range between 4.3 to 12 Hz in old mice was significantly lower than young mice (repeated measures AVOVA: factor Age F_(1,13)_=4.920, p=0.0450) (**Fig. 1d**). Since the activity of the theta frequency range during wakefulness is correlated with arousal^3,7^, old mice might have reduced arousal and less exploratory behavior compared to young mice. This result is consistent with the finding that old mice display an increased sleep propensity (**Fig. 1c**). The spectral power of the delta frequency range during NREM sleep is known as slow wave activity (SWA) and a good indicator of sleep intensity^7^. It has been reported that the absolute value of EEG SWA is significantly increased^3,4^ or tended to be increased^7^ in old mice. In our study, the power of the NREM EEG in the frequency range between 2.3-6.3 Hz in old mice was higher than young mice, but this trend did not reach statistical significance (repeated measures AVOVA: factor Age F_(1,13)_=2.029, p=0.1778) (**Fig. 1d**). The absolute value of SWA in old mice during a 24-hour period also tended to be increased compared to young mice (**Fig. 1e**). It has been suggested that absolute levels of SWA correlate with sleep pressure^3^, thus old mice might be exposed daily to a high sleep pressure compared with young mice. Although some studies showed significantly lower theta peak of REM EEG spectra in old mice, no notable differences were found in the REM EEG spectra in the frequency range between 4-9 Hz in our study (**Fig. 1d**).

### Old mice display increased sleep attempts during SD and homeostatic sleep response to SD

We next evaluated whether aging affects homeostatic sleep response by examining responses to SD in young and old mice. Sleep was disrupted by gentle handling for six hours, and then the mice were allowed to recover sleep loss (**Fig. 1f**). The number of sleep attempts gradually increased during SD in both young and old mice, and we noticed that old mice showed excessive sleepiness as their sleep attempts during SD were much greater than young mice (repeated measures ANOVA: factor Age F_(1,14)_=13.03, p=0.0028) (**Fig. 1g**). Thus, these results suggest that old mice might be more susceptible to accumulate sleep pressure from sleep loss than young mice. On the other hand, both young and old mice displayed a significant increase in SWA after SD (**Fig. 1h, Supplementary Fig. 1b**), indicating that the homeostatic response to SD is intact in old mice, which is consistent with other recent literature^3,4,7^. Surprisingly, the level of initial increase of SWA after SD in old mice was significantly higher than young mice (repeated measures ANOVA: factor Age x Time F_(5,70)_=3.583, p=0.0061), further supporting the notion that old mice might accumulate more sleep pressure during SD.

### The number of cFos+ cells in brain regions involved in the regulation of arousal and sleepiness increases significantly during SD

We examined which brain regions mainly responded to SD by staining the cFos protein, an immediate early gene product and a marker of neuronal activation^18^, in brain sections collected during SD, recovery sleep (RS), and control-sleep (SD-Cont and RS-Cont) (**Fig. 1f**). During SD, we found that the number of cFos+ cells was elevated in the brain regions known to regulate arousal and sleepiness, including the hypothalamus and brainstem. In the hypothalamus, the DMH, particularly at bregma -1.67 and -1.91mm, showed a greater number of cFos+ cells during SD compared to SD-Cont (**Fig. 2a-c, Supplementary Fig. 2a**). The median preoptic nucleus (MnPO) also showed increases in cFos+ cells during SD (**Supplementary Fig. 2b**), but not statistically significant. The number of cFos+ cells were significantly suppressed during RS in the DMH and MnPO compared with SD (**Fig. 2a, Supplementary Fig. 2b**). The LH, ventrolateral preoptic nucleus (VLPO) and tuberomammillary nucleus (TMN) exhibited no differences between SD-Cont and SD, although the number of cFos+ cells were significantly suppressed during RS in the LH and TMN compared with SD (**Supplementary Fig. 2b**). Therefore, neurons in the DMH and MnPO are activated specifically in response to sleep loss during SD. Whereas cFos+ cells in response to SD have been reported in the MnPO^19,20^, those in the DMH have been poorly characterized. Although DMH neurons are linked to aging and longevity control^12-14^ and also activated by psychological stress^21^, the involvement of DMH neurons in sleep control has not been fully elucidated. Thus, we decided to focus on these cFos+ cells in the DMH in response to SD.

**Fig. 2:**
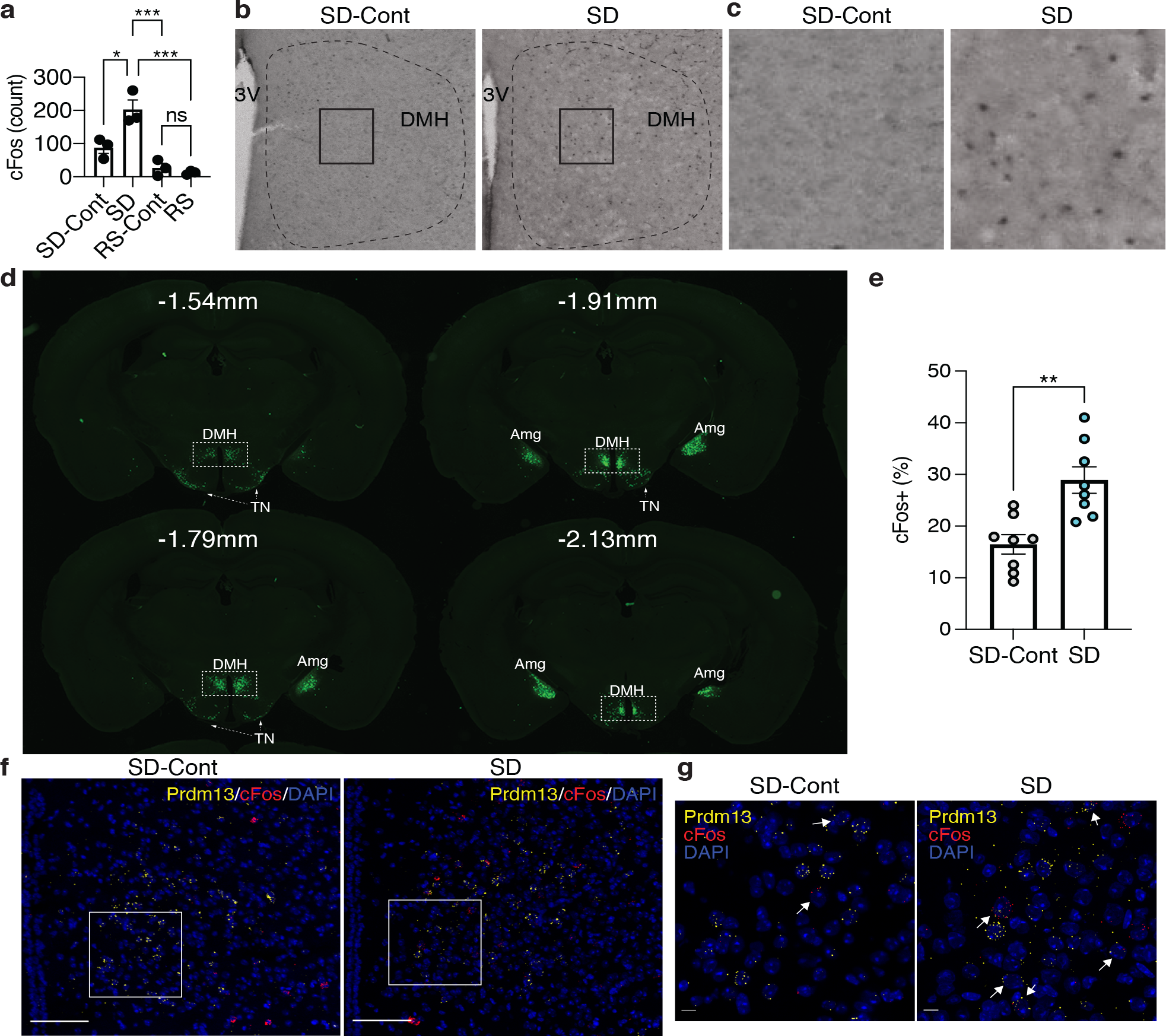
Prdm13+ neurons in the DMH are activated during SD. **a**, Numbers of cFos+ cells in the DMH during SD, RS and sleeping-control (SD-Cont, RS-Cont) detected by cFos immunohistochemistry. The total number of cFos+ cells in the DMH was counted at bregma -1.67 mm to, -1.79 mm and -1.91 mm and summed up (total three sections) each mouse (n=3). The third ventricle (3V) is shown. Values are shown as means ± S.E., *p<0.05, ***p<0.001, and non-significant (ns) by one-way ANOVA with Bonferroni’s post hoc test. **b**,**c**, Representative images of DMH sections at bregma -1.67 mm from mice under SD-Cont (left) and SD (right) with cFos. Boxed areas were shown at high magnification in **c. d**, Images of the ZsGreen signal including the DMH, amygdala (Amg) and tuberal nucleus (TN) at bregma -1.54, -1.79, -1.91 and -2.13 mm of *Prdm13*-ZsGreen mice. **e**, Ratios of *cFos*+ cells within *Prdm13*+ cells in young mice during SD and SD-Cont detected by RNAscope *in situ* hybridization (n=7-8). Values are shown as means ± S.E., *p<0.05 and **p<0.01 by two-way ANOVA with Bonferroni’s post hoc test. **f**,**g**, Representative images of DMH sections from young mice under SD-Cont (left) and SD (right) with *Prdm13* (yellow) and *cFos* (red) visualized by RNAscope. Boxed areas were shown at high magnification in **g**. Cells were counterstained with DAPI (blue). Scale bars indicate 100 and 10 μm (**f** and **g**, respectively).

### Prdm13+ neurons in the DMH are activated in response to SD

Given that *Prdm13* is one of the DMH-enriched genes and involved in sleep regulation^16^, we suspected that the cFos+ DMH cells responding to SD would include Prdm13+ neurons. To visualize Prdm13+ cells, *Prdm13*-CreERT2 mice were produced by targeted insertion of the coding sequence of tamoxifen-inducible Cre recombinase and 2A peptide into the native 3’ end of the *Prdm13* gene, generating the Prdm13-2A-CreERT2 protein. By crossing *Prdm13*-CreERT2 mice to Cre-dependent ZsGreen reporter mice, *Prdm13+* cells were visualized in the DMH and other brain regions such as the tuberal nucleus (TN) and amygdala (Amg) (**Fig. 2d**). No ZsGreen expression was observed without tamoxifen (data not shown). *In situ* hybridization confirmed ZsGreen+ cells were co-localized with endogenous *Prdm13* mRNA (**Supplementary Fig. 2c**). We also investigated electrophysiological characteristics of Prdm13+ DMH cells by whole cell patch-clamp technique. Using *Prdm13*-CreERT2-ZsGreen mice at 5-6 months of age, ZsGreen+ (Prdm13+) cells in the compact area of the DMH were selected (**Supplementary Fig. 2d**). We recorded synaptic activity (**Supplementary Fig. 2e**) and membrane capacitance (Cm), which is correlated with the morphology of neurons^22^ (**Supplementary Fig. 2f**), and confirmed that Prdm13+ DMH cells are electrically active cells, such as a neuron. Importantly, RNAscope analysis, a highly sensitive *in situ* hybridization method, revealed that the percentage of *cFos*+ cells among *Prdm13+* DMH neurons was significantly higher during SD than SD-Cont (**Fig. 2e-g**). Thus, Prdm13+ neuronal population in the DMH responds to sleep loss during SD.

### Mice with deficiency of *Prdm13* in the DMH display sleep fragmentation and excessive sleepiness during SD

To elucidate the role of Prdm13 signaling in age-associated sleep alterations, we generated DMH-*Prdm13*-KO mice. Our previous study demonstrated that *Prdm13* expression is partially regulated by Nkx2-1, which is highly expressed in the DMH^16^. We confirmed that most of Prdm13 is co-expressed with Nkx2-1 in the DMH, but not in the TN and Amg (**Supplementary Fig. 3a**). The percentage of Prdm13+Nkx2-1+ cells within Prdm13+ cells was 60±7.6%, 71±6.1% and 81±3.9% at bregma -1.43mm to -1.67mm, -1.79mm and -1.91mm, respectively (**Supplementary Fig. 3b**). Thus, we crossed *Prdm13-*floxed mice with *Nkx2-1*-CreERT2 mice to generate DMH-*Prdm13*-KO mice (**Fig. 3a**). The knockout efficiency of *Prdm13* in the DMH was about 70% after tamoxifen induction (**Fig. 3b**). Significant reduction of *Prdm13* expression was not observed in the TN and Amg of DMH-*Prdm13*-KO mice (**Fig. 3b**), and this event was specific to the hypothalamus since the expression of *Prdm13* remained intact in the retina where *Prdm13* is highly expressed (**Supplementary Fig. 3c**).

**Fig. 3:**
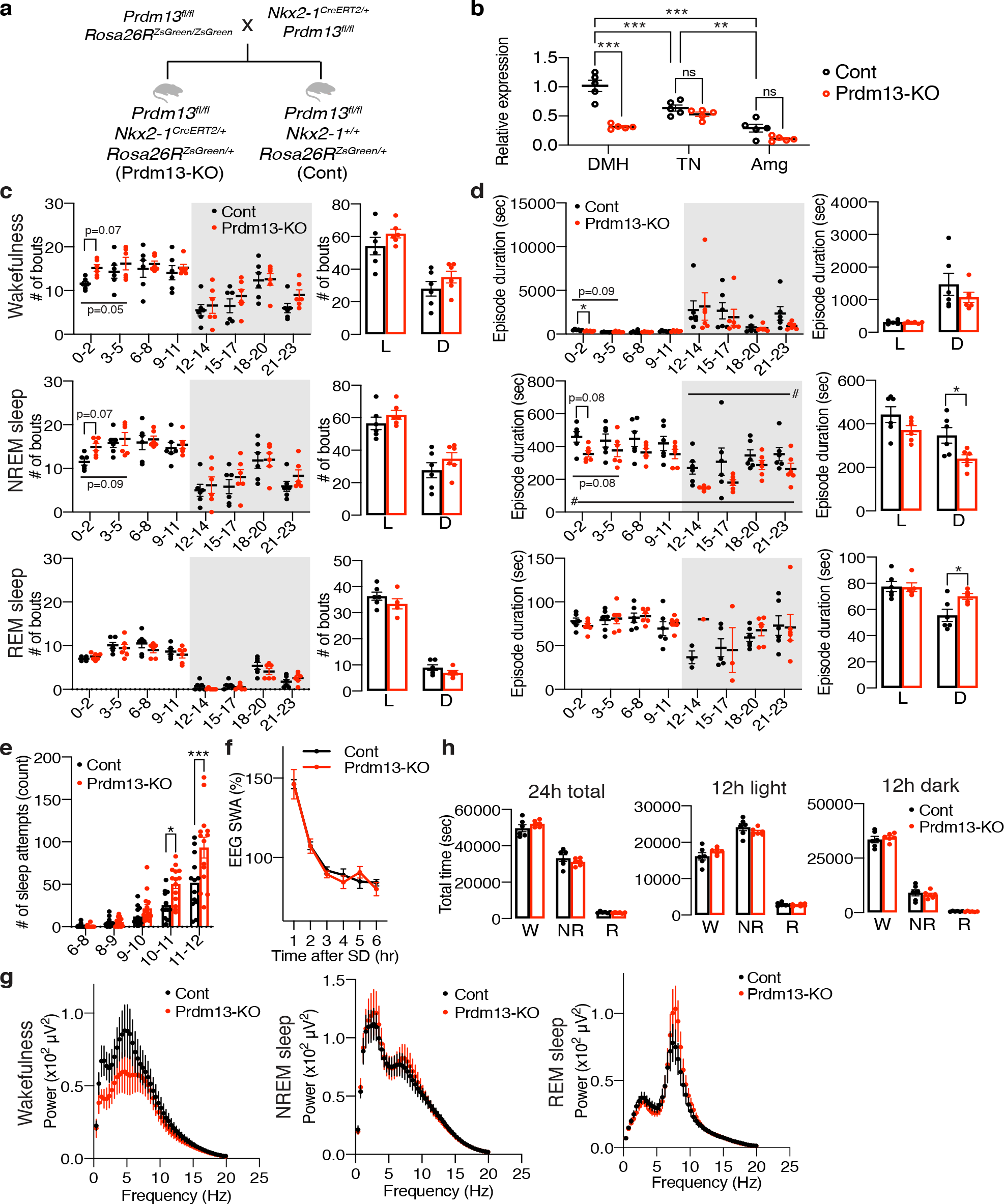
DMH-specific *Prdm13*-knockout mice display sleep alterations observed in aged C57BL/6J mice. **a**, Breeding strategy to generate DMH-specific *Prdm13*-knockout (Prdm13-KO) mice. After crossing *Prdm13*^*fl/f*^*;Rosa26R*^*ZsGreen/ZsGreen*^ mice and *Nkx2-1*^*CreERT2/+*^*;Prdm13*^*fl/fl*^ mice, *Prdm13*^*fl/fl*^*;Nkx2-1*^*CreERT2/+*^*;Rosa26R*^*ZsGreen/+*^ mice were used as Prdm13-KO mice and *Prdm13*^*fl/fl*^*;Nkx2-1*^*+/+*^*;Rosa26R*^*ZsGreen/+*^ mice were used as control (Cont) mice. **b**, Expression of *Prdm13* in the DMH, tuberal nucleus (TN) and amygdala (Amg) of Prdm13-KO and Cont mice (n=5). Values are shown as means ± S.E., **p<0.01, ***p<0.001 and non-significant (ns) by two-way ANOVA with Bonferroni’s post hoc test. **c**,**d**, Number of episodes (**c**) or duration (**d**) of wakefulness (top), NREM sleep (middle) and REM sleep (bottom) every 3 hours through a day (left) and during the light (L) and dark (D) periods (right) in Prdm13-KO and Cont mice. Shading indicates dark period (n=6). Values are shown as means ± S.E., #p<0.05 by repeated measures ANOVA, listed p-values and *p<0.05 by repeated measures ANOVA with Bonferroni’s post hoc test (left) or unpaired t-test (right). **e**, Number of sleep attempts during SD from 6am to 8am (6-8), 8am to 9am (8-9), 9am to 10am (9-10), 10am to 11am (10-11) and 11am to 12pm (11-12) in Prdm13-KO and Cont mice (n=13-14). Values are shown as means ± S.E., *p<0.05, ***p<0.001 by repeated measures ANOVA with Bonferroni’s post hoc test. **f**, SWA during NREM sleep after SD. Normalized power is relative to the average of the 24-hour baseline day each group (n=6). Values are shown as means ± S.E. **g**, Total amount of wakefulness, NREM sleep and REM sleep during a 24-hour period (24h total), 12-hour light period (12h light) or 12-hour dark period (12h dark) (n=6). Values are shown as means ± S.E. **h**, EEG spectra of wakefulness (left), NREM sleep (middle) and REM sleep (right) during the light period (n=5-6). Values are shown as means ± S.E.

We then analyzed sleep-wake patterns in DMH-*Prdm13*-KO and control mice at 4-6 months of age. During the light period between ZT0 to ZT5, DMH-*Prdm13*-KO mice showed a tendency of increase in the numbers of wakefulness and NREM sleep episodes compared with control mice (wakefulness; repeated measure ANOVA: factor Genotype F_(1,10)_=4.796, p=0.053, NREM sleep; repeated measure ANOVA: factor Genotype F_(1,10)_=3.539, p=0.089) (**Fig. 3c**). The duration of wakefulness episodes in DMH-*Prdm13*-KO mice was significantly shorter than control mice during the light period, and the duration of NREM sleep episodes in DMH-*Prdm13*-KO mice was significantly longer than control mice during the dark period (**Fig. 3d**). These results indicate that DMH-*Prdm13*-KO mice showed mild sleep fragmentation compared with control mice. We also assessed their responses to SD. The number of sleep attempts during SD in DMH-*Prdm13*-KO mice was significantly higher than those in control mice (repeated measure ANOVA: factor Genotype F_(1,25)_=9.131, p=0.0057) (**Fig. 3e**), recapitulating the phenotype of old wild-type mice (**Fig. 1g**). The level of initial increase of SWA after SD in DMH-*Prdm13*-KO mice was similar to control mice (**Fig. 3f, Supplementary Fig. 3d**), suggesting that the level of sleep pressure is comparable to each other. During wakefulness, the power of the EEG spectra at the frequency range between 4-12 Hz, in particular 4-9 Hz, in DMH-*Prdm13*-KO mice tended to be lower than control mice (repeated measures AVOVA: factor Genotype F_(1,9)_=0.7446, p=0.4106) (**Fig. 3g**), but there was no statistical significance. This trend was observed in old wild-type mice (**Fig. 1d**). The absolute value of SWA in DMH-*Prdm13*-KO mice during a 24-hour period tended to be higher compared to young mice (**Supplementary Fig. 3e**), but was not statistically significant. There were no abnormalities in the amounts of sleep and wakefulness, circadian period length and wheel-running activity in DMH-*Prdm13*-KO mice (**Fig. 3h, Supplementary Fig. 3f-h)**. Together, DMH-*Prdm13*-KO mice develop a moderate degree of sleep fragmentation, while the levels of sleep pressure and sleep propensity are still comparable with control mice at young age.

### Old DMH-*Prdm13*-KO mice display increased sleep fragmentation, adiposity, low physical activity and short lifespan compared to old control mice

To address the possibility that DMH-*Prdm13*-KO mice might accelerate physiological changes with advanced age, we conducted additional assessments using DMH-*Prdm13*-KO and control mice at 20 months of age. The level of sleep fragmentation in old DMH-*Prdm13*-KO mice was significantly higher than old control mice (**Fig. 4a,b**). In old DMH-*Prdm13*-KO mice, the number of wakefulness and NREM sleep episodes were significantly higher during the light period (**Fig. 4a**), and episode duration of wakefulness was significantly shorter (**Fig. 4b**) compared with old control mice. The duration of NREM sleep episodes in old DMH-*Prdm13*-KO mice was significantly shorter during the dark phase compared with old control mice (**Fig. 4b**). Therefore, long-term deficiency of Prdm13 signaling in the DMH worsens sleep fragmentation, particularly during the light period. The power of the EEG spectra at the frequency between 4-12 Hz, in particular 6.6 to 12 Hz, during wakefulness in old DMH-*Prdm13*-KO mice was significantly lower than old control mice (repeated measures AVOVA: factor Genotype F_(1,8)_=5.337, p=0.0497) (**Fig. 4c**), suggesting that old DMH-*Prdm13*-KO mice display increased sleep propensity compared with old control mice. No differences were found in the NREM and REM EEG spectra (**Fig. 4c**). The absolute value of SWA in old DMH-*Prdm13*-KO mice during a 24-hour period was tended to be higher compared to young mice (**Supplementary Fig. 4a**), but there was no statistical significance. Old DMH-*Prdm13*-KO mice displayed excessive sleepiness during SD compared with old controls (repeated measures ANOVA: factor Genotype F_(1,9)_=5.341, p=0.0462)(**Fig. 4d**). The level of initial increase of SWA after SD in old DMH-*Prdm13*-KO mice was significantly higher than old control mice (repeated measures ANOVA: factor Time x Genotype F_(5,45)_=5.024, p=0.0010) (**Fig. 4e, Supplementary Fig. 4b**). Therefore, old DMH-*Prdm13*-KO mice presumably accumulated more sleep pressure during SD compared to old control mice. The circadian period length and the amount of sleep and wakefulness were indistinguishable between old DMH-*Prdm13*-KO and control mice (**Supplementary Fig. 4c-e**), suggesting that circadian function, one of the major factors governing sleep-wake patterns^11^, was still intact in old DMH-*Prdm13*-KO mice. Although there was no change in body weight between DMH-*Prdm13*-KO and control mice at young age, DMH-*Prdm13*-KO mice gained more body weight than control mice at 18-20 months of age (**Fig. 4f**). The weight of perigonadal white adipose tissue in old DMH-*Prdm13*-KO mice tended to be higher than control mice (p=0.079 by unpaired t-test) (**Supplementary Fig. 4f**), and the size of adipocyte was significantly larger than control mice (**Supplementary Fig. 4g**,**h**). Moreover, the level of physical activity in old DMH-*Prdm13*-KO mice was significantly lower than old control mice (repeated measures ANOVA: factor Genotype F_(1,10)_=8.842, p=0.014) (**Fig. 4g**), while there was no change in food intake (**Supplementary Fig. 4i**). Taken together, DMH-*Prdm13*-KO mice exhibited the exacerbation of physiological decline with advanced age. Consistent with these observations, DMH-*Prdm13*-KO mice shortened their lifespan (p=0.0178 by log-rank test) (**Fig. 4h**). Since malignant neoplasm is the main cause of death in C57BL/6J mice^23^, we next tested whether Prdm13 deficiency in the DMH affects the incidence of malignant neoplasm. Most of the DMH-*Prdm13-*KO and control mice died by malignant neoplasm (83% and 86%, respectively), revealing that deletion of *Prdm13* in the DMH does not directly affect age-associated malignancy. Sleep alterations cause age-associated physiological dysfunctions^9,10,24^. Thus, alterations of sleep-wake patterns due to the deficiency of *Prdm13* may accelerate the decline in certain physiological function and reduce life expectancy without affecting age-associated malignancy.

**Fig. 4:**
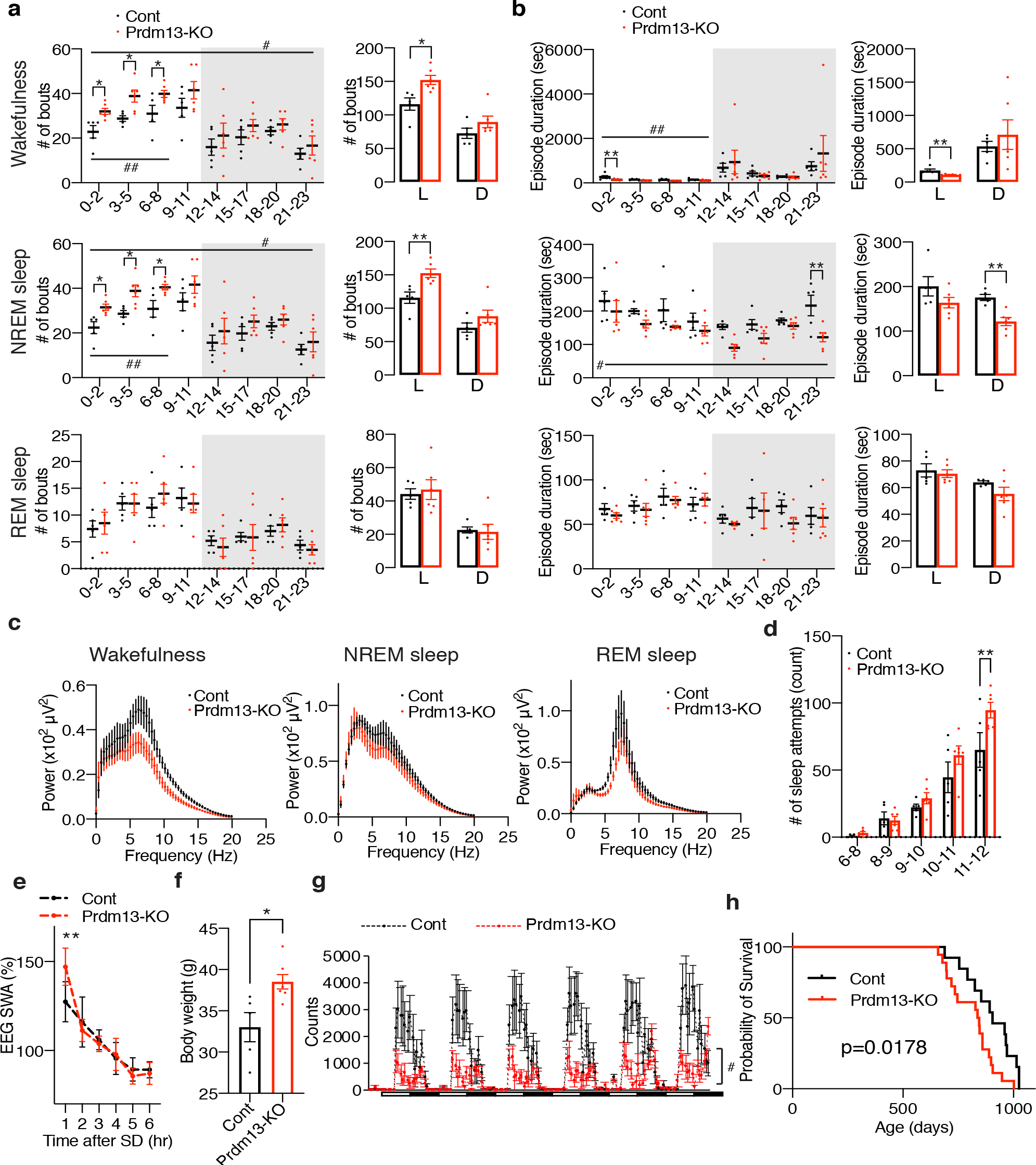
Old DMH-*Prdm13*-KO mice display age-associated pathophysiology and shortened lifespan. **a**,**b**, Numbers of episodes (**a**) and duration (**b**) of wakefulness (top), NREM sleep (middle) and REM sleep (bottom) every 3 hours through a day (left) and during the light (L) and dark (D) periods (right) in old DMH-specific *Prdm13*-knockout (Prdm13-KO) and control (Cont) mice (n=5-6). Values are shown as means ± S.E., #p<0.05 and ##p<0.01 by repeated measures ANOVA, *p<0.05 and **p<0.01 by repeated measures ANOVA with Bonferroni’s post hoc test (left) or unpaired t-test (right). **c**, EEG spectra of wakefulness (left), NREM sleep (middle) and REM sleep (right) during the light period (n=4-6). Values are shown as means ± S.E. **d**, Number of sleep attempts during SD from 6am to 8am (6-8), 8am to 9am (8-9), 9am to 10am (9-10), 10am to 11am (10-11) and 11am to 12pm (11-12) in old Prdm13-KO and Cont mice (n=5-6). Values are shown as means ± S.E., **p<0.01 by repeated measures ANOVA with Bonferroni’s post hoc test. **e**, SWA after SD of Prdm13-KO and Cont mice at 20 months of age. Normalized power is relative to the average of the 24-hour baseline day (n=5-6). Values are shown as means ± S.E., **p<0.01 by Bonferroni’s post hoc test. **f**, Body weight of old Prdm13-KO and Cont mice (n=5-7). Values are shown as means ± S.E., *p<0.05 by unpaired t-test. **g**, The level of wheel-running activity in old Prdm13-KO and Cont mice for six consecutive days (n=5-7). Values are shown as means ± S.E., #p<0.05 by repeated measures ANOVA. **h**, Kaplan-Meier curves of Prdm13-KO and Cont mice (n=13-18). Listed p-value was calculated by log-rank test.

### Short-term DR ameliorates age-associated sleep alterations in the presence of Prdm13 signaling

DR has been well known to ameliorate a wide variety of age-associated pathophysiological dysfunctions, delaying aging and extending lifespan. Thus, we speculated that DR could also ameliorate age-associated sleep alterations observed in old mice. After gradually decreasing the amount of food to 60% of daily food intake, mice at 20 months of age were kept under DR for 14 to 28 days (**Fig. 5a**). Control mice were fed *ad libitum* (AL). To minimize the disruption of their daily activity pattern, we fed mice at 5-6pm right before the dark period for both DR and AL mice. Every-day feeding did not cause disruption of the daily activity rhythm in AL and DR mice, except for some food-anticipatory activity^25,26^ observed right before lights-off in DR mice (**Supplementary Fig. 5a**), resulting in increased total amount of wakefulness in DR mice during the light period (**Supplementary Fig. 5c**). Body weights in old DR mice were significantly lower than old AL mice at 14, 21 and 28 days after dietary intervention (**Supplementary Fig. 5b**). Remarkably, the number of wakefulness and NREM sleep episodes in old DR mice were significantly lower than old AL mice during the light and dark periods, and the number of REM sleep episodes in old DR mice were significantly lower (**Fig. 5b**), whereas the durations of wakefulness, NREM and REM sleep episodes were longer than old-AL mice (**Fig. 5c**). Intriguingly, the power of EEG spectra across the overall frequency range during NREM and REM sleep in DR mice were significantly lower than AL mice (NREM sleep: repeated measures ANOVA: factor Diet F_(1,8)_=10.25, p=0.0126, REM: repeated measures ANOVA: factor Diet F_(1,8)_=5.405, p=0.0486) (**Fig. 5d**). It has been reported that starvation promotes significant reduction of EEG spectra due to hypothermia^27^. In fact, DR mice displayed significantly lower body temperature than AL mice over the course of experimental period (data not shown). Thus, decreases in EEG spectra in DR mice during NREM and REM sleep might be due to a low body temperature, not necessarily reflecting sleep pressure. The level of absolute SWA was significantly lower for a 24-hour period in DR mice but increased during a mealtime around ZT12 (**Supplementary Fig. 5d**), when body temperature is elevated, further supporting the idea that body temperature is associated with the regulation of EEG spectral power. In addition, DR significantly suppressed the number of sleep attempts during SD (repeated measures ANOVA: factor Diet F_(1,9)_=5.131, p=0.0498) (**Fig. 5e**). The level of SWA seen immediately after SD in DR mice was significantly lower than AL mice (repeated measures ANOVA: factor Time x Age F_(5,39)_=11.70, p<0.0001) (**Fig. 5f, Supplementary Fig. 5e**), suggesting that DR mice have less sleep pressure after SD compared with AL mice. Notably, SWA was increased in DR mice compared with AL mice at 6 hours after SD (ZT11) when SWA increased in the basal condition (**Supplementary Fig. 5d**), suggesting that daily rhythm of SWA is strongly persistent after DR. Taken together, DR effectively ameliorated age-associated sleep fragmentation and excessive sleepiness during SD.

**Fig. 5:**
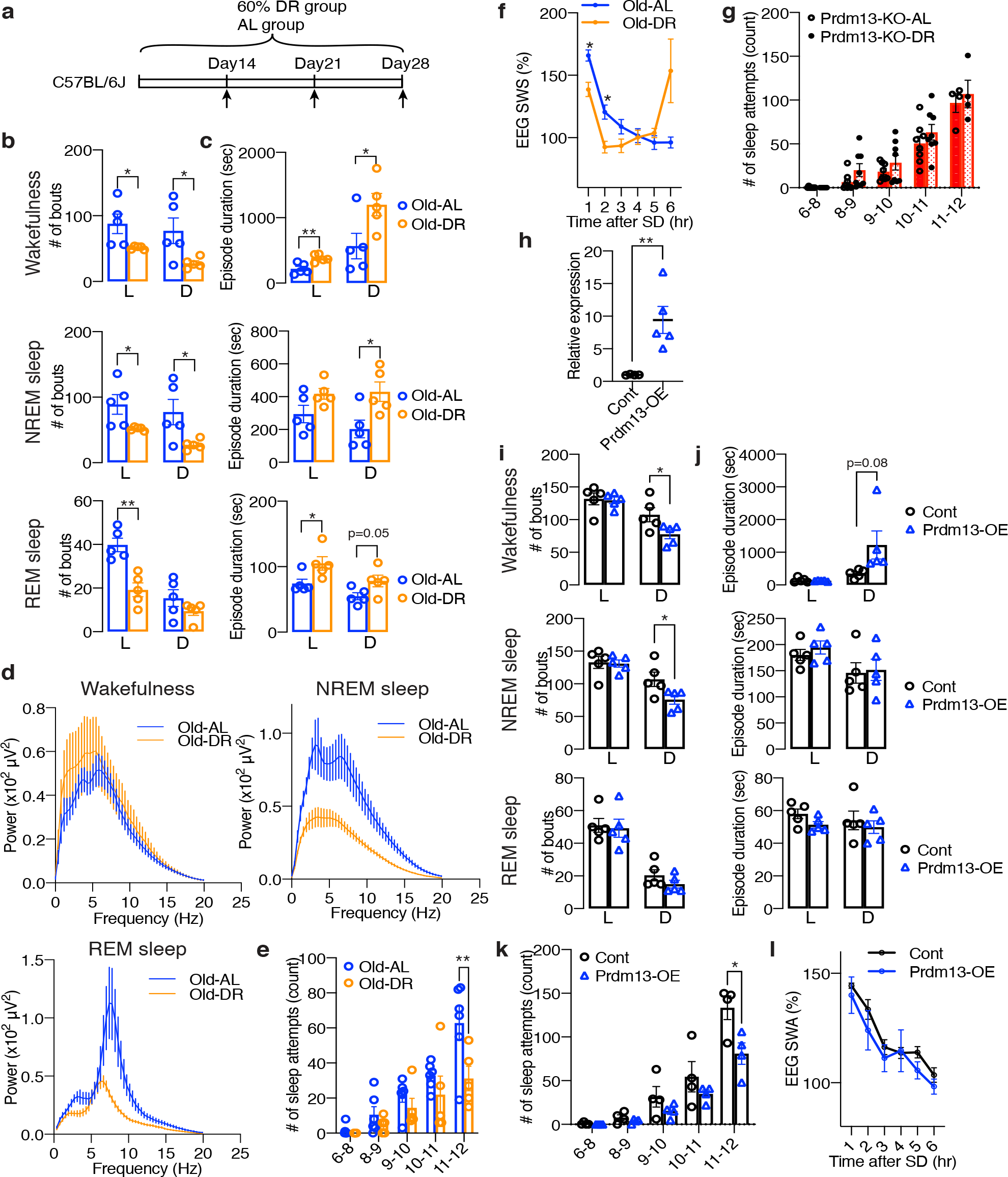
DR and overexpression of *Prdm13* in the DMH ameliorates age-associated sleep fragmentation and excessive sleepiness during SD. **a**, DR paradigm in C57BL/6J at 20 months of age. Mice at 20-months-old were fed under 60% diet or AL-diet for 14 to 28 days. **b**,**c**, Number of episodes (**b**) and duration (**c**) of wakefulness (top), NREM sleep (middle) and REM sleep (bottom) during the light (L) and dark (D) periods in AL and DR mice at 20 months of age (n=5). Values are shown as means ± S.E., listed p-value, *p<0.05 and **p<0.01 by unpaired t-test. **d**, EEG spectra of wakefulness (upper left), NREM sleep (upper right) and REM sleep (lower) during the light period (n=5). Values are shown as means ± S.E. **e**, Number of sleep attempts during SD from 6am to 8am (6-8), 8am to 9am (8-9), 9am to 10am (9-10), 10am to 11am (10-11) and 11am to 12pm (11-12) in AL and DR mice at 20 months of age (n=5-6). Values are shown as means ± S.E., **p<0.01 by repeated measures ANOVA with Bonferroni’s post hoc test. **f**, SWA after SD of AL and DR mice at 20 months of age. Normalized power is relative to the average of the 24-hour baseline day (n=5). Values are shown as means ± S.E., *p<0.05 by Bonferroni’s post hoc test. **g**, Number of sleep attempts during SD from 6am to 8am (6-8), 8am to 9am (8-9), 9am to 10am (9-10), 10am to 11am (10-11) and 11am to 12pm (11-12) in Prdm13-KO-AL and Prdm13-KO-DR mice (n=8). Values are shown as means ± S.E. **h**, Expression of *Prdm13* in the hypothalamus of *Prdm13*-overexpressing (Prdm13-OE) and control (Cont) mice (n=4-5). Values are shown as means ± S.E., *p<0.05 by unpaired t-test. **i**,**j**, Number of episodes (**i**) and duration (**j**) of wakefulness (top), NREM sleep (middle) and REM sleep (bottom) during the light (L) and dark (D) periods in Prdm13-OE and Cont mice (n=5). Values are shown as means ± S.E., *p<0.05 by unpaired t-test. **k**, Number of sleep attempts during SD from 6am to 8am (6-8), 8am to 9am (8-9), 9am to 10am (9-10), 10am to 11am (10-11) and 11am to 12pm (11-12) in Prdm13-OE and Cont mice (n=4). Values are shown as means ± S.E., *p<0.05 by repeated measures ANOVA with Bonferroni’s post hoc test. **l**, SWA after SD of Prdm13-OE and Cont mice. Normalized power is relative to the average of the 24-hour baseline day (n=4). Values are shown as means ± S.E.

Importantly, in DMH-*Prdm13*-KO mice that recapitulate the phenotypes of old wild-type mice, the effects of DR on sleep fragmentation and sleep attempts during SD were abrogated (**Fig. 5g, Supplementary Fig. 5f**,**g**). As expected, body weight was significantly lower in DMH-*Prdm13*-KO mice under DR compared with the same KO mice under AL-feeding, confirming that DR is properly conducted (**Supplementary Fig. 5h**). The number and durations of wakefulness and NREM sleep episodes in DMH-*Prdm13*-KO mice were indistinguishable between AL and DR (**Supplementary Fig. 5f**,**g**). On the other hand, the number of REM sleep episodes in DMH-*Prdm13*-KO under DR was significantly lower during the light period, but higher during the dark period than DMH-*Prdm13*-KO under AL (**Supplementary Fig. 5f**). In addition, the number of sleep attempts during SD was also indistinguishable between DR and AL in DMH-*Prdm13*-KO mice (repeated measures ANOVA: factor Diet F_(1,14)_=2.918, p=0.1097) (**Fig. 5g**). Except for food-anticipatory activity observed right before lights-off in DMH-*Prdm13*-KO-DR mice, the amount of sleep and wakefulness were indistinguishable between DMH-*Prdm13*-KO mice under DR and DMH-*Prdm13*-KO mice under AL (**Supplementary Fig. 5i**). Together, these results strongly suggest that Prdm13 is necessary to promote DR effects on sleep.

### Overexpression of *Prdm13* in the DMH ameliorates age-associated sleep alterations

Given that the level of hypothalamic *Prdm13*^16^ and its function decline with age, we next questioned whether overexpression of *Prdm13* in the DMH affects age-associated sleep alterations. We bilaterally injected lentivirus carrying a full-length *Prdm13* cDNA into the DMH of mice at 22 months of age. The level of *Prdm13* mRNA in DMH-specific *Prdm13-* overexpressing (*Prdm13-*OE) mice was 5 to 17-fold higher compared with control mice (**Fig. 5h**). Remarkably, the number of wakefulness and NREM sleep episodes in old *Prdm13-*OE mice were significantly lower, whereas duration of wakefulness in old *Prdm13-*OE mice was longer than old control mice, particularly during the dark period (**Fig. 5i,j**). Moreover, the number of sleep attempts during SD in old *Prdm13-*OE mice was significantly lower than control mice (**Fig. 5k**). The level of SWA after SD in *Prdm13-*OE mice did not differ from old control mice (**Fig. 5l, Supplementary Fig. 5j**). Thus, the restoration of Prdm13 signaling in the DMH partially rescues age-associated sleep alterations.

### Prdm13 functions as a transcription factor in the DMH

What is the molecular function of Prdm13 in the DMH? We found that the DMH expresses previously uncharacterized alternative splicing variants of *Prdm13* by 5’-rapid amplification of cDNA-ends (RACE)-PCR analysis (**Supplementary Fig. 6a**,**b**). To further characterize the function of hypothalamic Prdm13 (htPrdm13), we developed an antibody against this variant. This antibody specifically detected the recombinant and overexpressed htPrdm13 proteins (data not shown) and also the deletion of Prdm13 in the DMH-*Prdm13*-KO mice (**Fig. 6a**), confirming its specificity. Using this antibody, we examined the subcellular localization of the Prdm13 protein by biochemical fractionation (**Fig. 6b left**). The Prdm13 protein was found exclusively in the RNase- and DNase-resistant nuclear scaffold fraction from wild-type hypothalami (**Fig. 6b right**). Nuclear localization of Prdm13 in the hypothalamus was also confirmed by using hypothalami from a newly developed mouse model expressing podoplanin (PA)-tagged Prdm13 (**Fig. 6c**). These results indicate that Prdm13 likely functions as a transcription factor in the DMH. This is consistent with a previous report showing that Prdm13, reported as Prdm13-202, acts as a transcription factor in the dorsal neural tube^28^. Intriguingly, among DMH-enriched genes that were previously reported^16^, the levels of *cholecystokinin (Cck), gastrin releasing peptide (Grp)*, and *pro-melanin-concentrating hormone (Pmch)* mRNA were significantly reduced in the compact region of the DMH (Bregma -1.79 and -1.91) of DMH-*Prdm13*-KO mice (**Fig. 6d**). Transcriptional activities of *Cck* (one-way ANOVA, F_(1.016, 3.047)_=10.89, p=0.0446), *Grp* (one-way ANOVA, F_(2.158, 6.474)_=11.63, p=0.0068), and *Pmch* (one-way ANOVA, F_(1.547, 4.641)_=54.50, p=0.0007) promoters were upregulated by Prdm13-202 in a dose-dependent manner (**Fig. 6e**). As well as Prdm13-202, htPrdm13 also upregulated *Cck* transcription (one-way ANOVA, F_(1.010, 2.020)_=49.88, p=0.0189) (**Fig. 6f**). Moreover, a Prdm13-zinc-finger (Zif) mutant with amino acid mutations (C187A, H207A, C622A, H638A, C650A, H666A, C679A, H695A), leading to inactivation of four Zif domains, showed significantly decreased transcriptional activity, whereas a Prdm13-PR/SET deletion mutant still activated these promoters to levels similar to Prdm13-202 (**Fig. 6e**). These results reveal that the Zif domain, but not the PR/SET domain, is necessary for Prdm13 to upregulate the activity of *Cck, Grp* and *Pmch* promoters.

**Fig. 6:**
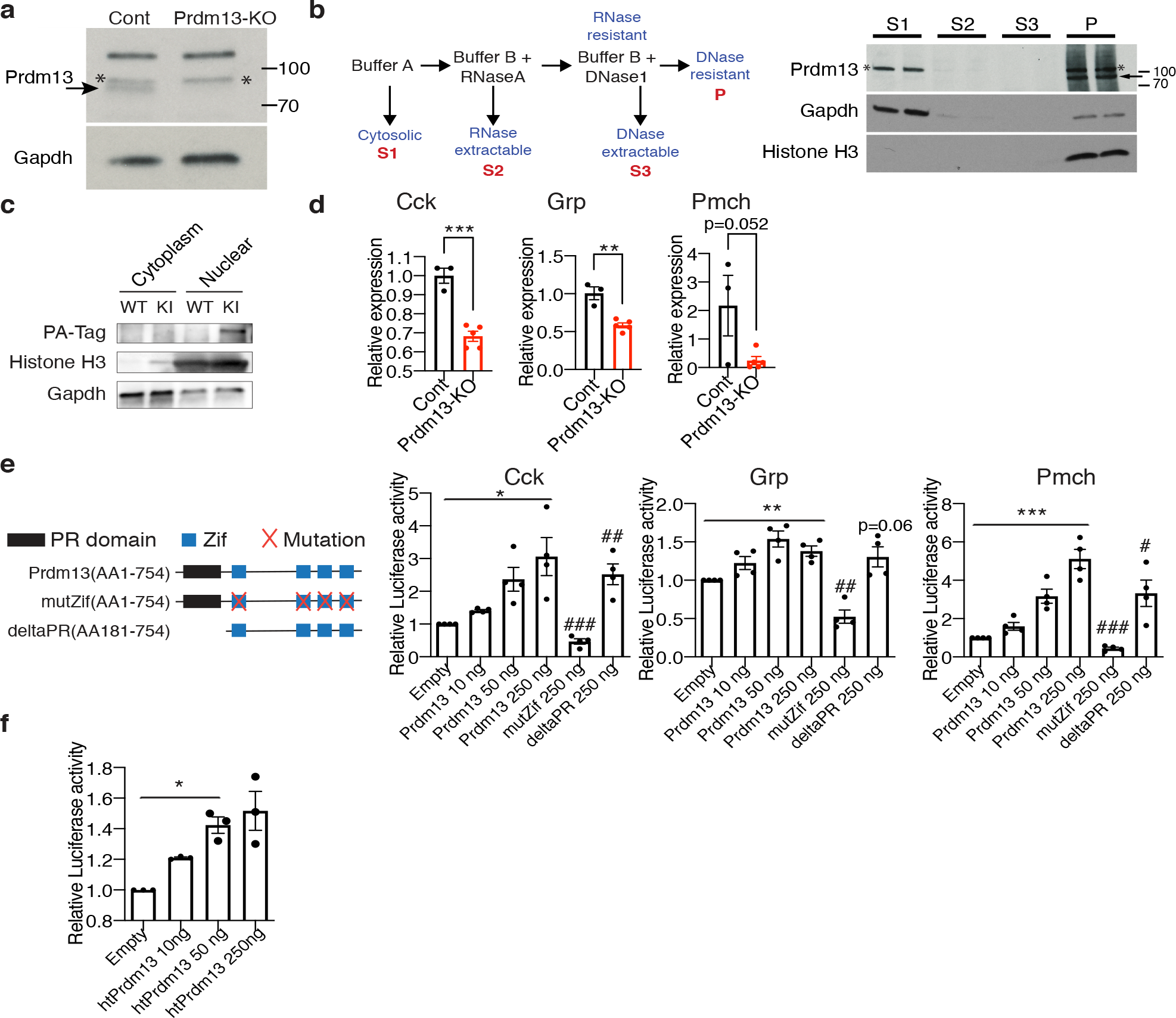
Prdm13 in the DMH is a transcription factor. **a**, Western blot of Prdm13 in DMH collected by laser microdissection from DMH-specific Prdm13-KO and Cont mice (n=4 mice/lane). The arrow indicates the band for Prdm13; asterisks (*) indicate non-specific bands. **b**, Schematic of fractionation protocol from mouse hypothalami (left). Western blot of Prdm13 in hypothalamic fractions of C57BL/6J mice (right). Hypothalami from two C57BL/6J female mice were combined for each lane, and 8% equivalent of each fraction was run on the gel. Cytoplasmic supernatant (S1), RNase-extractable supernatant (S2), DNase-extractable supernatant (S3), and insoluble pellet (P) were run each lane. The arrow indicates the band for Prdm13; asterisks (*) indicate non-specific bands. **c**, Western blot of Prdm13 in hypothalamic fractions of *Prdm13*-PA-Tag (KI) and wild-type (WT) mice. Cytosolic and nuclear fractions were run each lane as indicated. **d**, Expression of *Cck, Grp* and *Pmch* mRNA in the DMH of DMH-*Prdm13*-KO (Prdm13-KO) and control (Cont) mice (n=3-5). Values are shown as means ± S.E., listed p-value, **p<0.01 and ***p<0.001 by unpaired t-test. **e**, Transcriptional activity of Prdm13-202 and Prdm13-mutants for the luciferase reporter vector containing the promoter region of *Cck, Grp* and *Pmch*. Schematic representation of Prdm13-202 and Prdm13-mutants are shown above. NIH3T3 cells were co-transfected with 250 ng of luciferase reporter plasmid and plasmid expressing Prdm13-202 (Prdm13), Prdm13-Zif mutant (mutZif) or Prdm13-deltaPR mutant (deltaPR). Obtained luminescence was normalized to total protein concentration (n=3, four individual experiments). Values are shown as means ± S.E., listed p-value, *p<0.05, **p<0.01 and ***p<0.001 by one-way ANOVA with Bonferroni’s post hoc test, ^#^p<0.05 and ^##^p<0.01 and ^###^p<0.001 by unpaired t-test. **f**, Transcriptional activity of hypothalamic Prdm13 (htPrdm13) for the luciferase reporter plasmid containing the promoter region of *Cck*. NIH3T3 cells were co-transfected with 250 ng of reporter plasmid and 10, 50 or 250 ng of *htPrdm13*-expressing plasmid. Obtained luminescence was normalized to total protein concentrations (n=3, three individual experiments). Values are shown as means ± S.E., *p<0.05 by one-way ANOVA with Bonferroni’s post hoc test.

### DMH Prdm13+Cck+ neurons were activated during SD

Notably, at bregma -1.67 mm in the DMH, *Prdm13* was co-localized with *Cck* or *Grp* within the *Prdm13*+ neuronal population about 26% and 14%, respectively, (**Fig. 7a,b**), but showed almost no co-localization with *Pmch* (**Supplementary Fig. 7a**). The percentage of *Prdm13+Cck+* in *Prdm13+* DMH neurons was significantly higher than the percentage of *Prdm13+Grp+* neurons (**Fig. 7b**). Therefore, *Prdm13* might functionally or mechanistically connect with *Cck* and/or *Grp* in the DMH. *Cck*+ cells among *Prdm13+* DMH neurons, particularly at bregma -1.67 mm, were widely distributed, but significantly more predominant in the medial part than the lateral part (16.1±1.3% and 9.8±1.3% in the medial and lateral parts, respectively, unpaired t-test: p<0.001), while *Grp*+ cells among *Prdm13+* DMH neurons were distributed mainly in the lateral part (2.9±1.1% and 10.9±1.3% in the medial and lateral parts, respectively, unpaired t-test: p<0.001) (**Fig. 7a,b**). During SD, 58% of *Prdm13+cFos+* neurons in the DMH were localized in the medial part (**Fig. 7c,d**), where *Prdm13+Cck+* neurons were their majority, whereas only a few *Prdm13+Grp+* neurons were observed (**Supplementary Fig. 7b**). Thus, we questioned whether Cck is involved in the response to SD in this particular area. The percentage of *cFos+* cells within *Prdm13*+*Cck+* DMH neuronal population during SD was significantly higher than those in SD-Cont in young mice (**Fig. 7e right**), but not *Prdm13*+*Cck-* DMH neuronal population (**Fig. 7e left**). Thus, the specific neuronal population expressing both Prdm13 and Cck is activated in response to SD. Unexpectedly, the percentage of *cFos+* cells within *Prdm13*+*Cck+* DMH neuronal population during SD in old mice was also significantly higher than those in SD-Cont (**Fig. 7f**). However, the degree of increase in cFos+ in response to SD in young mice (2.0-fold) noticeably dropped in old mice (1.5-fold). Furthermore, the level of *Cck* in the hypothalamus of old mice was significantly lower compared with young mice (**Supplementary Fig. 7c**), and the number of *Prdm13*+ DMH cells that highly expressed Cck in old mice tended to be lower than young mice (**Supplementary Fig. 7d)**. Therefore, decreased level of Cck expression in the DMH might affect sleep-wake patterns in old mice, and these effects might occur with affecting neuronal activation and another neuronal event.

**Fig. 7:**
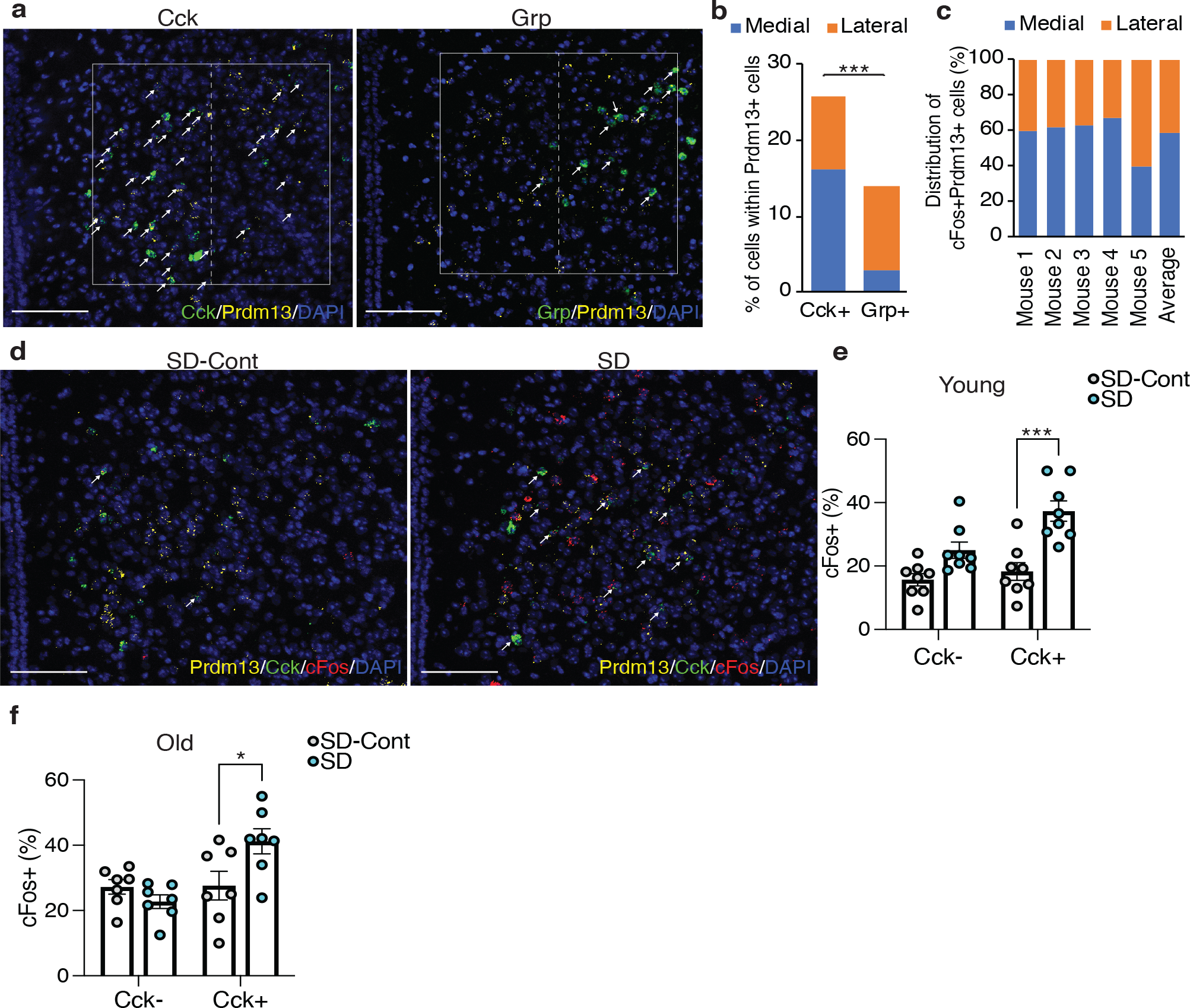
DMH Prdm13+Cck+ neurons are activated during SD. **a**, Representative images of the DMH with *Prdm13* (yellow) and one of the two genes, *Cck* or *Grp* (green) visualized by RNAscope. Cells were counterstained with DAPI (blue). White boxes show the DMH, which is divided into medial and lateral areas by dashed lines. White arrows show yellow+green+ cells. Scale bar indicates 100 μm. **b**, Ratios of *Cck*+ or *Grp*+ cells within *Prdm13*+ cells in medial, lateral or total (medial and lateral) DMH (n=3-5). Values are shown as means ± S.E., ***p<0.001 by unpaired t-test. **c**, Distribution of *cFos*+*Prdm13*+ cells (n=5). **d**, Representative images of the DMH from young mice under SD-Cont and SD with *Prdm13* (yellow), *Cck* (green) and *cFos* (red) visualized by RNAscope. Cells were counterstained with DAPI (blue). White arrows show *Prdm13+Cck+cFos+* cells. Scale bar indicates 100 μm. **e**,**f**, Ratios of *cFos*+ cells within *Prdm13*+*Cck*-(left) or *Prdm13*+*Cck*+ (right) cells in young (**e**) or old (**f**) mice during SD-Cont and SD (n=7-8). Values are shown as means ± S.E., *p<0.05, ***p<0.001 by two-way ANOVA with Bonferroni’s post hoc test.

## Discussion

We demonstrated that Prdm13 signaling in the DMH is responsible for the remarkable effect of DR against age-associated sleep fragmentation and excessive sleepiness during SD in mice. Consistently, in humans, two years of 25% DR promoted better sleep quality reflected by lower scores reported on the Pittsburgh Sleep Quality Index compared with the AL group^29^. Another study showed that two days of 10% DR significantly increased the duration of stage 4 deep NREM sleep^30^. Our study revealed that the deficiency of Prdm13 signaling promotes phenotypes similar to age-associated sleep-wake alterations, but overexpression of *Prdm13* in the DMH of old mice mitigates those age-associated sleep changes. Thus, enhancing Prdm13 signaling in the DMH might be beneficial not only to prevent age-associated sleep alterations, but also to ameliorate age-associated pathophysiologies in old animals. Further elucidating detailed downstream molecular events regulated by Prdm13 signaling and characteristics of Prdm13+ DMH neurons will be of great interest to explore a potential intervention on age-associated sleep-wake patterns.

A potential mechanism by which Prdm13 signaling in the DMH controls sleep fragmentation and excessive sleepiness during SD is the transcriptional regulation of neuropeptides in the DMH. In particular, *Cck* transcription is downregulated within the hypothalamus of old mice. Optogenetic and chemogenetic studies show that activation of GABAergic/Cck+ neurons in the preoptic area of the hypothalamus promotes NREM sleep^31^. Similarly, the activation of glutamatergic/Cck+ neurons in the perioculomotor region of the midbrain also promotes NREM sleep likely through the activation of GABAergic neurons in the preoptic area of the hypothalamus. Together, it is likely that activation of Cck+ neurons in the preoptic area of the hypothalamus and the perioculomotor region of the midbrain promotes sleep. Young DMH-*Prdm13*-KO mice display excessive sleepiness during SD to the extent equivalent to old mice. Because sleep fragmentation is further developed with age, it is conceivable that the decreased function of Prdm13 signaling causes excessive sleepiness during SD, and then sleep fragmentation. The detailed mechanisms by which the deficiency of *Prdm13* leads to sleep fragmentation still need to be elucidated in future studies.

At young age, DMH-*Prdm13*-KO mice display sleep fragmentation and excessive sleepiness during SD, mimicking sleep changes observed in old wild-type mice. In addition, DMH-*Prdm13*-KO mice show exacerbated sleep fragmentation and develop other physiological changes such as increased adiposity and decreased wheel-running activity over age. The reason for DMH-*Prdm13*-KO mice to show changes primarily in sleep, and secondarily in body weight and physical activity at older age is currently unknown. One possibility is that prolonged sleep fragmentation induces low physical activity and increased adiposity. To support this idea, it has been reported that 20 days of sleep fragmentation during the light period promotes a decreased physical activity in young mice^9^. In humans, low sleep efficiency, a widely recognized index of sleep consolidation and fragmentation, is significantly associated with the reduction in daytime physical activity^32,33^. Therefore, the low physical activity observed in old DMH-*Prdm13*-KO mice might be a consequence of chronic sleep fragmentation. Similar to physical activity, obesity is also promoted by sleep fragmentation through an increased food intake in mice^10,24^, but old DMH-*Prdm13*-KO mice do not alter their food intake. Therefore, the increased adiposity in old DMH-*Prdm13*-KO mice might be a consequence of low physical activity.

In this study, we examined age-associated changes in sleep-wake patterns, EEG spectra and responsiveness to SD. Consistent with previous reports^2-4,7^, we confirmed that old mice showed reduced total amount of wakefulness and increased NREM sleep during a 24-hour period. Such differences are more pronounced during the dark period, but we also observed differences during the light period. Such differences could be due to a sexual dimorphism in sleep-wake patterns^34^ because our study mainly used female mice for sleep analysis, whereas other studies have used male mice^2-4,7^. Nonetheless, our findings in this study implicate an important possibility that old mice are more susceptible to accumulate sleep pressure compared with young mice. In fact, the level of SWA after SD, which reflects accumulated sleep pressure from increased wakefulness, in old mice was significantly higher than young mice. Indeed, our finding is consistent with a notion that old mice live under a high sleep pressure^3^. Since the homeostatic sleep response is fairly intact with age, accumulation and generation of sleep pressure in old mice might be greater than in young mice. However, there are clear discrepancies between mice and humans in sleep studies^35^. It should be noted that some of the age-associated sleep alterations such as sleep fragmentation are conserved in both mice and human, whereas basic sleep architecture is quite different between each other (e.g., polyphasic vs. monophasic sleep, nocturnal vs. diurnal). In most reports, aged mice show an increase in total NREM sleep in the dark period^2-4,7^, but older humans show a decrease^3,4^. In this regard, it is interesting that a most recent study has reported that a recessive mutation to the human PRDM13 gene causes ataxia with cerebellar hypoplasia and delayed puberty with hypogonadotropic hypogonadism^36^. Recent GWAS study revealed that PRDM13 is listed as one of the potential nocturnal enuresis risk genes^37^. It remains unclear whether these patients also have sleep defects. Thus, while mice are useful for modeling certain aspects of aging and sleep, an extra caution will be necessary in extrapolating the results obtained from mice to humans.

We discovered Prdm13 signaling in the DMH affects sleep-wake patterns during the aging process. We also uncovered that Prdm13 signaling is necessary to promote DR effects in age-associated sleep patterns and restoration of Prdm13 in the DMH ameliorates age-associated sleep alterations. Our study also elucidated that Prdm13 acts as a transcription factor that regulates the expression of critical neuropeptides in the DMH. Among those neuropeptides, *Cck* is most likely involved in the regulation of age-associated sleep-wake pattern changes mediated by Prdm13 functioning as a transcription factor in the DMH. Other downstream target genes of Prdm13 may also potentially be involved in the age-associated regulation of sleep-wake patterns. The detailed relationship between the activity of Prdm13+ DMH neurons and Prdm13 signaling itself still need to be elucidated in the future studies.

## Acknowledgements

A.S. is supported by grants JSPS KAKENHI (17H07417, JP18H03186, 20K21780), the Japan Agency for Medical Research and Development (AMED) (JP20gm5010001s0604), the Sleep Medicine Foundation, the Takeda Science Foundation, and the Research Fund for Longevity Sciences from the NCGG (28–47). S.T. (1st author) is supported by a grant from Suntory Foundation for Life Science. M.W. and N.R. are supported by a grant NIH P50 HD103525 to the Washington University Intellectual and Developmental Disabilities Research Center. S.I. is supported by grants from the National Institute on Aging (AG037457, AG047902) and the Tanaka Fund at Washington University School of Medicine and also by AMED (JP20gm5010002s0003) at the IBRI. K.N. is supported by grants from AMED (JP20gm5010002s0304) and JST Moonshot R&D (JPMJMS2023).

## Contributions

Conceptualization, S.T., C.S.B., R.Y., Y.T., S.I. and A.S.; Methodology, S.T., C.S.B., R.Y., Y.T., H.T., N.R., S.M., J.A., Y.K., K.N., N.O., S.T., S.T., S.I. and A.S.; Investigation, S.T., R.Y. and A.S. conducted most of experiments, C.S.B. conducted experiments necessary for Fig. 3b,6a,6b,6d, H.T. conducted experiments necessary for Supplementary Fig. 2d-f; Writing-Original Draft, S.T. and A.S.; Writing-Review & Editing; C.S.B., N.R., K.N., M.W. and S.I. All other authors also contributed to the final manuscript.

## Methods

### Animal models

All mouse experiments and procedures were approved by the Animal Care and Use of the NCGG. Mice were housed in 12/12-hour light/dark cycle (lights on at 6am and off at 6pm) with free access to food and water. 1-2 month-old C57BL/6J mice were purchased from the Charles River Laboratories International, Inc. (Yokohama, Japan), and grew up to 4-6 or 18-20 months of age (as young and old groups, respectively) in our Animal Facility at the NCGG and aged mice specialized suits at the NCGG. *Rosa26R*^*ZsGreen/ZsGreen*^ and *Nkx2-1*^*CreERT2/+*^mice (Jackson stock no: 007906 and 014552) were obtained from Jackson Laboratory. *Prdm13*^*fl/fl*^ mice (RIKEN BRC stock no: RBRC09371)^38^ were obtained from RIKEN BRC. For DR study, C57BL/6J mice at 20 or 4 months of age or DMH-*Prdm13*-KO mice at 12 months of age were fed under 60% diet or AL-diet for 28 days. To minimize habitual stress and disruption of daily pattern, food was gradually decreased to 60% of daily food intake, and both AL and DR groups were fed daily at 5-6pm right before the dark period. Mice were closely monitored and included daily body weight measurement during the experimental period. Female mice were mainly used for sleep studies, wheel-running analysis, food intake behavior studies, and DR studies. Both male and female mice were used to confirm excessive sleepiness during SD in DMH-*Prdm13*-KO mice. Only male mice were used in longevity study and the DR study using DMH-*Prdm13*-KO mice.

*Prdm13*-CreERT2 mice were generated by the Laboratory Animal Resource Center at the University of Tsukuba. The detailed procedure was described previously^39^. Briefly, a targeting vector was designed to insert the *Prdm13* sgRNAs (5’
s-GAC TCC TAA CGC GCC TTC CA-3’) into pX330-mC plasmid, which carried Cas9-mC expression unit^39^ (pX330-mC-Prdm13sgRNA). pCreERT2-Prdm13 was designed to insert a 2A peptide (P2A), CreER^T2^ recombinase, and rabbit globin polyadenylation signal, replacing TAA stop codon in the fourth exon of *Prdm13* gene. These two constructs (pX330-mC-Prdm13sgRNA and pCreERT2-Prdm13) were microinjected into zygotes from C57BL/6J mice. Subsequently, injected zygotes were transferred into oviducts in pseudopregnant ICR female mice (Charles River Laboratories International, Inc. Yokohama, Japan), and 85 newborns were obtained. The designed knock-in (KI) mutation was confirmed by PCR using the following primers: Prdm13 screening 5Fw: 5’-CAT GCA CAG CAC TTG TGG TAG AGA AAT C-3’, Prdm13 screening 3Rv: 5’-ATT TAG AAT TGG AGC AAA CAG GGG GAT T-3’. No random integrations were detected by PCR with primers detecting the ampicillin resistance gene.

*Prdm13*-PA-Tag KI mice were generated by the Laboratory Animal Resource Center at the University of Tsukuba. We attempted to introduce the PA-Tag coding sequence connected to the LG3-linker sequence just before the TAA stop codon of *Prdm13* gene^40^. Briefly, the gRNA (5’
s-AGT CCC TGG AAG GCG CGT T-3’) was synthesized and purified by GeneArt™ Precision gRNA Synthesis Kit (Thermo Fisher Scientific). In addition, we designed a 200-nt single-stranded DNA oligonucleotide (ssODN) donor, placing the LG3-PA sequence between the genomic regions from 54 bp upstream of the TAA stop codon to 53 bp downstream of the TAA (Integrated DNA Technologies). The gRNA, ssODN, and GeneArt™ Platinum™ Cas9 Nuclease (Thermo Fisher Scientific) were electroporated to C57BL/6J zygotes using a NEPA 21 electroporator (NEPAGNENE), as described previously^41^. After electroporation, 2-cell embryos were transferred into oviducts in pseudopregnant ICR female mice, and 32 newborns were obtained. The PA-Tag KI mutation was confirmed by PCR using the following primers: Prdm13 QC primer F: 5’-TCA ACA AGC ACA TCC GAC TC-3’, Prdm13 QC primer R: 5’-TGA CGT GAT CCT GAA CCT CA-3’. The PCR products were sequenced by using BigDye Terminator v3.1 Cycle Sequencing Kit (Thermo Fisher Scientific).

### Sleep analysis

Isoflurane-anesthetized mice were surgically implanted with stainless screw electrodes placed over the right frontal bone for reference and right/left parietal bone for active recording electroencephalogram (EEG), and wire electrodes in the nuchal muscle for electromyogram (EMG) recording. All signals were grounded to a bone screw electrode placed over the cerebellum midline. Mice were recovered from surgery for three days and subsequently acclimatized to the recording cage for three weeks. EEG/EMG recording was performed continuously for 2 consecutive days. Recording electrodes were connected to TBSI Tethered System T8 amplifier (TBSI) via T8 Headstage (A50-2139-G3, TBSI), a lightweight cable and commutator to enable free movement and feeding in a sound and light proof enclosure with a 12/12-hour light/dark cycle. EEG/EMG signals were digitized at 600 Hz, filtered at 0.3-35 Hz for EEG and 10-100 Hz for EMG by PowerLab system (ADInstruments). Wireless EEG Logger (ELG-2, Bio research Center) was used for sleep analysis in Fig. 5i-l. 10-second epochs of EEG/EMG signals were semiautomatically scored as wakefulness, NREM sleep, and REM sleep by SleepSign (KISSEI COMTEC) with visual examination. Score was blinded for genotypes during quantification. Spectra analysis was performed by a FFT (FFT; 0.4-20 Hz, 0.38 Hz resolution). Three outliers were detected using Grubb’s and ROFU tests (Graph Pad Prizm 9) (one young mice in Fig. 1, one KO mouse in Fig. 3, 4) and excluded from spectrum analysis. SWA during NREM sleep was computed across the 24-hour recording period by SleepSign (KISSEI COMTEC). SWA after SD was normalized to the average of SWA for the average of 24-hour period each mouse.

### SD study

Mice were individually housed prior to the experiment. On the day of SD, food was removed at 6am and mice were kept awake using a long Q-tip until 12pm (6 hours SD) by gentle touching of mice, as previously reported^42^. Attempts to sleep were determined by the onset of behaviors typical of sleep such as cornering, curling, and eye closing. Once such behaviors were observed, we placed the long Q-tip in front of the mouse. One sleep attempt was counted when the mouse did not react to it. EEG/EMG recording was performed during SD to monitor the effectiveness of the SD protocol. All animals indicated <5% of sleep during the six hours of SD. After SD, food was added, and the mice were allowed to sleep. Genotypes and conditions were blinded during the experimental procedures.

### Immunohistochemistry and immunofluorescence

Mice were anesthetized with isoflurane and perfused with PBS followed by 4% paraformaldehyde (PFA) at 11am for SD and SD-Cont, and at 2pm for RS and RS-Cont. Brains were fixed with 4% PFA overnight and placed into 30% sucrose until saturated.

Thirty-micrometer cryosections were collected into PBS and stored in cryoprotectant at −20°C. For immunofluorescent staining, samples were stained using primary antibodies: anti-Nkx2-1 (TTF-1) (1:500, ab76013, Abcam) and secondary antibodies. To stain cFos, samples were stained with anti-cFos (1:1,000, 226003, Synaptic Systems) and universal biotinylated anti-mouse/rabbit IgG (Universal Elite ABC kit, PK-7200, Vector laboratories) antibodies with Universal Elite ABC kit and developed with Vector SG Substrate Kit, Peroxidase (SK-4700, Vector laboratories). The number of cFos-positive cells was quantified by visual scoring. Genotypes and conditions were blinded during the experimental procedures.

### *In situ* hybridization

For RNAscope, brains from C57BL/6J mice were dissected, embedded in OCT and frozen on dry ice. The embedded frozen blocks were cut at 14 *μ*m thick using cryostat CM1850 (Leica) and mounted on slides. The sections were stored at -80°C until further processing. Target mRNA was detected using the RNAscope Multiplex Fluorescent Reagent Kit v2 [Advanced Cell Diagnostics (ACD)]. RNA probes (ACD) used in this study are as follows: *Prdm13* (Cat# 543551-C2), *Fos* (Cat# 316921-C3), *Cck* (Cat# 402271), *Grp* (Cat# 317861), and *Pmch* (Cat# 478721). The frozen sections were fixed in pre-chilled 4% PFA in PBS for 10 min. After 2 times washing with PBS, the sections were dehydrated through 50%, 70%, 100% and 100% ethanol for 5 min each. The slides were air dried for 5min. The slides were treated with hydrogen peroxide for 10 min at room temperature. Probe hybridization and signal amplification were performed using TSA Plus kit (PerkinElmer) according to the ACD’s instructions. The slides were counter stained with DAPI and mounted using 2.5% 1,4,diazabicyclo[2.2.2]octane (DABCO) in 50% glycerol. The slides were imaged with LSM700 laser-scanning confocal microscope (Zeiss) with ZEN 2009 software (Zeiss). Cells positive for *Prdm13, cFos, Cck, Grp* and *Pmch* were manually detected. To evaluate *Cck* expression level semiquantitatively, signal dots derived from *Cck* mRNA were counted manually and categorized into 4 grades: 1 (1-5 dots/ cell), 2 (6-10 dots/ cell), 3 (11-15 dots/ cell) and 4 (>16 dots/ cell). Genotypes and conditions were blinded during the experimental procedures.

### Lentivirus production

To generate the *Prdm13*-expressing lentiviral construct, *Prdm13* cDNA was cloned into the FCIV.FM1 vector (a gift from the Viral Vectors Core at Washington University School of Medicine). High-tittered viruses were generated from the Viral Vectors Core at Washington University School of Medicine. Briefly, lentiviruses were produced by co-transfecting HEK293T cells with the *Prdm13*-expressing vectors and three packaging vectors (pMD-Lg, pCMV-G, and RSV-REV) by the calcium phosphate precipitation procedure. Six hours after transfection, the medium was replaced with the complete medium containing 6 mM sodium butyrate. Culture supernatant was collected 42 hours after transfection. The supernatant was passed through a 0.45 *μ*m filter, concentrated by ultracentrifugation through a 20% sucrose cushion, and stored at -80°C until use. Virus titer was determined by transducing HT1080 cells and assaying for reporter expression using flow cytometry.

### Lentivirus injections

Following anesthesia with isoflurane gas, the mouse was placed in a three-point fixation stereotactic frame. Bregma was identified, and appropriate coordinates for the stereotactic injection were registered: relative to Bregma for the DMH, anterior-posterior (AP) -1.4 mm, medial-lateral (ML) ± 0.3 mm, and dorsal ventral (DV) -5.4 mm. A burr hole was made using a dental drill, and a glass capillary was directed to the previously determined coordinates. Viruses were slowly injected (100 nL/ 2 min). After the injection, animals were allowed to recover in a temperature regulated incubator (32°C) until fully awake. All injected mice had four weeks to fully recover before being used for any experiments. Viruses with the following titers and volumes were injected: Lentiviruses carrying *Prdm13* or *fLuc* cDNA (2.0 × 10^8^ IU/mL, 500 nL in the DMH).

### Whole cell patch-clamp electrophysiology

Mice were anesthetized with an isoflurane–oxygen mixture, and the brain was removed. The brain was quickly transferred into ice-cold dissection buffer (25 mM NaHCO_3_, 1.25 mM NaH_2_PO_4_, 2.5 mM KCl, 0.5 mM CaCl_2_, 7 mM MgCl_2_, 25 mM glucose, 110 mM choline chloride, 11.6 mM ascorbic acid, 3.1 mM pyruvic acid and 1mM kynurenic acid), gassed with 5%CO_2_/95%O_2_. Coronal brain slices were cut (300 µm; Leica VT1200S) in dissection buffer. The slices were then incubated in physiological solution (118 mM NaCl, 2.5 mM KCl, 26 mM NaHCO_3_, 1 mM NaH_2_PO_4_, 10 mM glucose, 4 mM MgCl_2_, 4 mM CaCl_2_, pH 7.4, gassed with 5%CO_2_/95%O_2_).

Patch recording pipettes (3–7 MΩ) were filled with intracellular solution (115 mM cesium methanesulfonate, 20 mM CsCl, 10 mM HEPES, 2.5 mM MgCl_2_, 4 mM Na_2_ATP, 0.4 mM Na_3_GTP, 10 mM sodium phosphocreatine and 0.6 mM EGTA at pH 7.25). To record the mEPSC (−60 mV holding potential) or mIPSC (0 mV holding potential), the recording chamber was perfused with physiological solution with 0.5 µM TTX. Whole-cell recordings were obtained from Prdm13+ neurons of the DMH with a Multiclamp 700B (Axon Instruments). Cells with membrane resistance > 100 MΩ and series resistance < 20 MΩ were only recorded. Whole-cell patch-clamp data were collected with Clampex and analyzed using Clampfit 10.7 software (Axon Instruments)^43^.

### Identification of 5’-end of htPrdm13 cDNA from mouse hypothalami

We performed 5’-rapid amplification of cDNA-ends (5’-RACE)-PCR analyses to determine the 5’ end of htPrdm13 transcripts in RNAs isolated from the hypothalamus of C57BL/6J. The hypothalamus of C57BL/6J were dissected and immediately frozen in liquid nitrogen. Total RNA from the hypothalamus was extracted with RNeasy kit (QIAGEN). 5’-RACE was performed by SMARTer RACE 5’/3’ kit (Clontech) by manufacture protocol. Briefly, first-strand cDNA was synthesized with 5’ RACE CDE Primer A (Clontech) (5’-RACE-Ready cDNA samples), and the 5’-RACE-Ready cDNA sample was used for 5’-RACE reaction with a htPrdm13 specific primer (5’-GATTACGCCAAGCTT-TAGCGAAAGGTCCTCCAGCAGTA-3’). RACE products were purified by NuclearSpin Gel (QIAGEN) and PCR Clean-Up Kit (QIAGEN). Purified RACE products were inserted into pRACE vector with In-Fusion DM Master Mix. 5’-sequencing was confirmed by reading more than seven independent clones in 5’-RACE products using primer (5’-AAGCTTGGCGTAATC-3’). PCR was conducted using 5’-end primer (5’-ATGGTGAGAGGGGAGCTGGT-3’) and 3’-end primer (5’-TTAGGAGTCGTGCTCGCCAC-3’) to confirm amplification of htPrdm13 using hypothalamic cDNAs.

### Western blot analysis of Prdm13 from mouse hypothalamus

For antibody confirmation, the compact region of DMH from four DMH-*Prdm13*-KO or control mice was collected by laser microdissection into Laemmli’s sample buffer using the Leica LMD 6000 system (Leica) and boiled 5 min. The detailed procedure for sample preparation was described previously^44^. For fractionation, two C57BL/6J mouse hypothalami were dissected into Buffer A [10 mM HEPES-KOH, pH 7.9, 10 mM KCl, 1.5 mM MgCl_2_, 0.5 mM DTT, 1 mM PMSF, protease inhibitor cocktail (Roche), 1 mM NaF, 1 mM Na_3_VO_4_]. The tissue was incubated on ice to swell 20 min. Samples were homogenized 10 sec on medium speed using a Polytron homogenizer and centrifuged 3 min at 4,000 rpm. The resulting cytoplasmic supernatant (S1) was removed. The remaining pellet was washed with Buffer A and centrifuged again. The washed pellet was resuspended in Buffer B (20 mM HEPES-KOH, pH 7.9, 400 mM NaCl, 1.5 mM MgCl_2_, 200 mM EDTA, 0.5 mM DTT, 1 mM PMSF, protease inhibitor cocktail, 1 mM NaF, 1 mM Na_3_VO_4_) with 200 *μ*g/mL RNaseA and the sample was rocked 30 min at room temp and centrifuged 3 min at 14,000 rpm. The resulting RNase-extractable supernatant (S2) was removed. The remaining pellet was washed with Buffer B and centrifuged again. The washed pellet was resuspended in Buffer B with 300 *μ*g/ml DNase I (QIAGEN) and 5 mM MgCl_2_. The sample was incubated 30 min at 37°C, mixing periodically, then centrifuged 3 min at 14,000 rpm. The resulting DNase-extractable supernatant (S3) was removed. The remaining pellet was washed with Buffer B and centrifuged again. 2x Laemmli’s sample buffer was added to the insoluble pellet (P), homogenized with syringe and 28G needle, and boiled 5 min. After centrifugation, no pellet remained. To make protein extracts from S1-S3 fractions, 5x Laemmli’s sample buffer was added and samples were boiled 5 min. The 8% equivalent of each fraction by volume was run on a 4-15% TGX gel (Bio-Rad) for Western blotting using affinity-purified polyclonal rabbit anti-mouse Prdm13. Antibodies for Western blotting included affinity-purified polyclonal rabbit anti-mouse htPrdm13 (Covance), anti-Gapdh antibody (MAB374MI, Thermo Fisher Scientific) and anti-Histone H3 antibody (#9715, Cell Signaling Technology).

### Western blot analysis of Prdm13 using Prdm13-PA-Tag hypothalami

Hypothalami from *Prdm13*-PA-Tag KI C57BL/6J mice or wild-type C57BL6J mice were dissected and frozen in liquid nitrogen. One mouse brain was homogenized using a syringe and needle in 60 *μ*L of lysis buffer A (10 mM HEPES, pH 7.5, 2 mM MgCl_2_, 3 mM CaCl_2_, 300 mM sucrose, 1 mM DTT) supplemented with Halt Protease and Phosphatase Inhibitor Cocktail (Thermo Fisher Scientific). The homogenates were incubated 10 min on ice and centrifuged at 600xg for 5 min. The supernatants were transferred to new tubes as cytoplasmic fractions. The pellets were resuspended in 60 *μ*L of lysis buffer B (50 mM HEPES, pH 7.4, 150 mM NaCl, 2.5% SDS, 2 mM MgCl_2_, 1 mM DTT) supplemented with Halt Protease and Phosphatase Inhibitor Cocktail and homogenized using a syringe and needle. The homogenates were centrifuged at 16,000xg for 20 min. The supernatants were transferred to new tubes as nuclear fractions. Protein concentration was determined using the BCA protein assay kit (Takara Bio).

Equal amounts of protein extracts were resolved by SDS–PAGE using 4-15% Mini-PROTEAN TGX precast gel (Bio-Rad) and transferred to PVDF membrane. Prdm13-PA was detected using anti-PA-Tag antibody conjugated with horse radish peroxidase (HRP) (015-25951, Fujifirm). Histone H3 and Gapdh were detected using anti-histone H3 antibody (#9715, Cell Signaling Technology) or anti-Gapdh antibody (MA5-15738, Thermo Fisher Scientific) as primary antibodies, and anti-rabbit IgG HRP linked whole antibody (NA934V, GE Healthcare) or anti-mouse IgG HRP linked whole antibody (NA931V, GE Healthcare) as secondary antibodies. Protein bands were visualized using Amersham ECL select (Cytiva).

### Plasmid construction for reporter assay

Hypothalamic *Prdm13*-coding sequence was amplified from a cDNA of C57BL/6J mouse hypothalamus using primers containing a FLAG-tag-coding sequence. The amplified DNA fragment was introduced into pcDNA3.1(+) mammalian expression vector (Thermo Fisher Scientific) using *Eco*RI and *Xba*I sites, creating pcDNA3.1-Prdm13 plasmid, which expresses C-terminal FLAG-tagged hypothalamic *Prdm13* driven by the CMV promoter. A plasmid expressing FLAG-tagged Prdm13-202 or mutants of Prdm13-202 was constructed as follows. The coding sequence of amino acids (AA) 99-754 of Prdm13-202 and C-terminal FLAG tag was PCR amplified from the pcDNA3.1-Prdm13 plasmid. The amplified DNA fragment and a DNA fragment coding AA 1-98 of Prdm13-202 synthesized as a gBlocks Gene Fragment (Integrated DNA Technologies), were assembled into FCIV.FM1 vector using NEBuilder HiFi DNA Assembly Master Mix (New England Biolabs), creating a plasmid expressing C-terminal FLAG-tagged Prdm13-202 driven by a ubiquitin promoter. Similarly, the coding sequence of AA 181-754 of Prdm13-202 was amplified and introduced into FCIV.FM1, creating a plasmid expressing the PR domain-deletion mutant of Prdm13-202, named Prdm13 deltaPR. Prdm13-202 with amino acid mutations (C187A, H207A, C622A, H638A, C650A, H666A, C679A, H695A) leading to inactivation of four zinc finger domains^28^ named Prdm13 mutZif. A plasmid expressing Prdm13 mutZif was created by introducing DNA fragment coding Prdm13 mutZif synthesized as a gBlocks Gene Fragment (Integrated DNA Technologies) into FCIV.FM1 vector. Luciferase reporter vectors were constructed as follows. Approximately 4 kbp upstream sequence from the transcription start site of mouse *Cck, Grp* or *Pmch* gene was PCR amplified and inserted into pGL4.1 (Promega) using *Kpn*I and *Hin*dIII sites.

### Gene expression analysis of DMH samples

The compact region of the DMH was collected by laser microdissection using the Leica LMD 6000 system (Leica). The detailed procedure for sample preparation was described previously^44^. Total RNA was extracted following laser microdissection using the PicoPure RNA isolation kit (Applied Biosystems). cDNA was synthesized using the Applied Biosystems High Capacity cDNA Reverse Transcription Kit. Quantitative real-time RT-PCR was conducted, and relative expression levels were calculated for each gene by normalizing to *Gapdh* levels and then to the average of the control samples. Primers used in this study were Mm99999915_g1 (*Gapdh*), Mm00446170_m1 (*Cck*), Mm00612977_m1 (Grp), Mm01242886_g1 (*Pmch*) and Mm01217509_m1 (*Prdm13*) (Applied Biosystems).

### Reporter assay for *Prdm13* transcriptional variants and mutants

250 ng of the reporter plasmid and 250 ng of the expression plasmid were transiently co-transfected into NIH3T3 cells^45^ (a gift from Dr. Sugimoto) using HilyMax transfection reagent (Dojindo). For mock assay, 250 ng of empty FCIV.FM1 vector was used instead of the expression vector. After 24 hours, luminescence was measured using the dual luciferase reporter system (Promega) without detecting renilla luciferase activity. Obtained luminescence was normalized to total protein concentration measured by BCA protein assay kit (Takara Bio).

### Wheel-running analysis

Mice were individually housed into cage with the Wireless Running Wheel (Med Associates Inc.), and habituated for two weeks. Basal physical activity was recorded with Wheel Manager Software (Med Associates Inc.) for 4-5 days under 12/12-hour light/dark cycle. After the basal measurement, physical activity recorded under constant darkness for 10 days. Physical activity and period length were determined by Wheel Analysis Software (Med Associates Inc.). Light/dark-cycles were strictly monitored by a light censor (Brain Science Idea Co. Ltd.).

### Measurement of adipocyte size

Mice were anesthetized with isoflurane and perfused with PBS followed by 4% PFA. The perigonadal WAT were fixed with 4% PFA overnight and placed into 70% ethanol. Paraffin sections were prepared by Tissue Tech VIRTM 5 Junior (VIP-5-Jr-10, Sakura Fine Chemical), and HE-staining was conducted by Multiple Slide Stainer (DRS 2000-B, Sakura Fine Chemical). Slide images were scanned by Nanozoomer (Hamamatsu Photonics). The sections were viewed at 20x magnification and randomly selected five areas each section using Nanozoomer NDP.view2 (Hamamatsu Photonics). The size of adipocytes was measured by ImageJ with Adipocyte_Tools.ijm. Genotypes and conditions were blinded during the experimental procedures.

### Longevity study

All animals were kept in our animal facility with free access to standard laboratory diet and water. No mice used for the longevity study were used for any other biochemical, physiological, or metabolic tests. The endpoint of life was the time when each mouse was found dead during daily inspection. Moribund mice were euthanized according to our institutional animal care guidelines, and the time at euthanasia was its endpoint. Survival data of each cohort were analyzed by plotting the Kaplan-Meier curve and performing the log-rank test using Prizm. Tumor and organ tissues were dissected immediately after the animals were euthanized or death, fixed with 10% formalin neutral buffer solution (Fujifilm) for 24 hours, and processed for paraffin-embedded sections, followed by hematoxylin-eosin staining for pathological diagnosis. Immunostainings were performed, if necessary, for differentiation of tumors, by BOND MAX/III (Leica) with BOND Polymer Refine Detection (ds9800; Leica). Antibodies against CD68 (1:2,000 dilution; histiocytic marker; E3O7V; #97778; Cell Signaling Technology), alfa-fetoprotein (1:500 dilution; 14550-1-AP; Proteintech), CD45R (1:300 dilution; B-cell marker; B220; #550286; BD Biosciences; with F[ab’]2 anti-rat IgG[H&L]; 1:500 dilution; 712-4126; Rockland antibodies& assays) and CD3 (1:500 dilution; T-cell marker; 21120-1-AP; Proteintech) were used.

### Statistical Analysis

Excel or Graph Pad Prizm 9 software was used for data quantification and generation of graph. One-way ANOVA followed by Bonferroni’s post hoc test was employed for comparisons between three or more groups. Repeated measures ANOVA followed by Bonferroni’s post hoc test was employed for number of bouts or episode duration every 3 hours for a 24-hour period, EEG SWA for a 24-hour period and after SD, number of sleep attempts during SD, amount of wakefulness, NREM sleep and REM sleep for a 24-hour period and physical activity. Two-way ANOVA followed by Bonferroni’s post hoc test was employed for testing the differences between the age groups and experimental conditions. FFT significance was determined by two-way ANOVA. Number of bouts and episode duration significance at each period (light or dark period), total amount of wakefulness, NREM or REM sleep each period (light, dark, or total 24-hour period) were determined by unpaired Student’s *t-*test. Pierson’s correlation was employed for examining the relationship between number of bouts and number of sleep attempts, or between total number of sleep attempts and remaining lifespan. Log-rank test was employed for longevity study.

## References

1 Foley, D. J. et al. Sleep complaints among elderly persons: an epidemiologic study of three communities. Sleep 18, 425–432, doi:10.1093/sleep/18.6.425 (1995).

2 Soltani, S. et al. Sleep-Wake Cycle in Young and Older Mice. Front Syst Neurosci 13, 51, doi:10.3389/fnsys.2019.00051 (2019).

3 Panagiotou, M., Vyazovskiy, V. V., Meijer, J. H. & Deboer, T. Differences in electroencephalographic non-rapid-eye movement sleep slow-wave characteristics between young and old mice. Sci Rep 7, 43656, doi:10.1038/srep43656 (2017).

4 McKillop, L. E. et al. Effects of Aging on Cortical Neural Dynamics and Local Sleep Homeostasis in Mice. J Neurosci 38, 3911–3928, doi:10.1523/JNEUROSCI.2513-17.2018 (2018).

5 Mander, B. A., Winer, J. R. & Walker, M. P. Sleep and Human Aging. Neuron 94, 19–36, doi:10.1016/j.neuron.2017.02.004 (2017).

6 Carskadon, M. A., Brown, E. D. & Dement, W. C. Sleep fragmentation in the elderly: relationship to daytime sleep tendency. Neurobiol Aging 3, 321–327, doi:10.1016/0197-4580(82)90020-3 (1982).

7 Wimmer, M. E. et al. Aging in mice reduces the ability to sustain sleep/wake states. PLoS One 8, e81880, doi:10.1371/journal.pone.0081880 (2013).

8 Naidoo, N., Ferber, M., Master, M., Zhu, Y. & Pack, A. I. Aging impairs the unfolded protein response to sleep deprivation and leads to proapoptotic signaling. J Neurosci 28, 6539–6548, doi:10.1523/JNEUROSCI.5685-07.2008 (2008).

9 He, J., Kastin, A. J., Wang, Y. & Pan, W. Sleep fragmentation has differential effects on obese and lean mice. J Mol Neurosci 55, 644–652, doi:10.1007/s12031-014-0403-7 (2015).

10 Hakim, F. et al. Chronic sleep fragmentation during the sleep period induces hypothalamic endoplasmic reticulum stress and PTP1b-mediated leptin resistance in male mice. Sleep 38, 31–40, doi:10.5665/sleep.4320 (2015).

11 Scammell, T. E., Arrigoni, E. & Lipton, J. O. Neural Circuitry of Wakefulness and Sleep. Neuron 93, 747–765, doi:10.1016/j.neuron.2017.01.014 (2017).

12 Zhang, G. et al. Hypothalamic programming of systemic ageing involving IKK-beta, NF-kappaB and GnRH. Nature 497, 211–216, doi:10.1038/nature12143 (2013).

13 Satoh, A. et al. Sirt1 extends life span and delays aging in mice through the regulation of Nk2 homeobox 1 in the DMH and LH. Cell Metab 18, 416–430, doi:10.1016/j.cmet.2013.07.013 (2013).

14 Zhang, Y. et al. Hypothalamic stem cells control ageing speed partly through exosomal miRNAs. Nature 548, 52–57, doi:10.1038/nature23282 (2017).

15 Yoshida, M. et al. Extracellular Vesicle-Contained eNAMPT Delays Aging and Extends Lifespan in Mice. Cell Metab 30, 329–342 e325, doi:10.1016/j.cmet.2019.05.015 (2019).

16 Satoh, A., Brace, C. S., Rensing, N. & Imai, S. Deficiency of Prdm13, a dorsomedial hypothalamus-enriched gene, mimics age-associated changes in sleep quality and adiposity. Aging Cell 14, 209–218, doi:10.1111/acel.12299 (2015).

17 Fontana, L., Partridge, L. & Longo, V. D. Extending healthy life span--from yeast to humans. Science 328, 321–326, doi:10.1126/science.1172539 (2010).

18 Gao, Y. J. & Ji, R. R. c-Fos and pERK, which is a better marker for neuronal activation and central sensitization after noxious stimulation and tissue injury? Open Pain J 2, 11–17, doi:10.2174/1876386300902010011 (2009).

19 Brown, R. E., Basheer, R., McKenna, J. T., Strecker, R. E. & McCarley, R. W. Control of sleep and wakefulness. Physiol Rev 92, 1087–1187, doi:10.1152/physrev.00032.2011 (2012).

20 Alam, M. A., Kumar, S., McGinty, D., Alam, M. N. & Szymusiak, R. Neuronal activity in the preoptic hypothalamus during sleep deprivation and recovery sleep. J Neurophysiol 111, 287–299, doi:10.1152/jn.00504.2013 (2014).

21 Kataoka, N., Hioki, H., Kaneko, T. & Nakamura, K. Psychological stress activates a dorsomedial hypothalamus-medullary raphe circuit driving brown adipose tissue thermogenesis and hyperthermia. Cell Metab 20, 346–358, doi:10.1016/j.cmet.2014.05.018 (2014).

22 Liu, Y. B., Lio, P. A., Pasternak, J. F. & Trommer, B. L. Developmental changes in membrane properties and postsynaptic currents of granule cells in rat dentate gyrus. J Neurophysiol 76, 1074–1088, doi:10.1152/jn.1996.76.2.1074 (1996).

23 Blackwell, B. N., Bucci, T. J., Hart, R. W. & Turturro, A. Longevity, body weight, and neoplasia in ad libitum-fed and diet-restricted C57BL6 mice fed NIH-31 open formula diet. Toxicol Pathol 23, 570–582, doi:10.1177/019262339502300503 (1995).

24 Wang, Y. et al. Chronic sleep fragmentation promotes obesity in young adult mice. Obesity (Silver Spring) 22, 758–762, doi:10.1002/oby.20616 (2014).

25 Stephan, F. K. The “other” circadian system: food as a Zeitgeber. J Biol Rhythms 17, 284–292, doi:10.1177/074873040201700402 (2002).

26 Pendergast, J. S. & Yamazaki, S. The Mysterious Food-Entrainable Oscillator: Insights from Mutant and Engineered Mouse Models. J Biol Rhythms 33, 458–474, doi:10.1177/0748730418789043 (2018).

27 Huang, Y. G. et al. The relationship between fasting-induced torpor, sleep, and wakefulness in laboratory mice. Sleep 44, doi:10.1093/sleep/zsab093 (2021).

28 Chang, J. C. et al. Prdm13 mediates the balance of inhibitory and excitatory neurons in somatosensory circuits. Dev Cell 25, 182–195, doi:10.1016/j.devcel.2013.02.015 (2013).

29 Martin, C. K. et al. Effect of Calorie Restriction on Mood, Quality of Life, Sleep, and Sexual Function in Healthy Nonobese Adults: The CALERIE 2 Randomized Clinical Trial. JAMA Intern Med 176, 743–752, doi:10.1001/jamainternmed.2016.1189 (2016).

30 Collet, T. H. et al. The Sleep/Wake Cycle is Directly Modulated by Changes in Energy Balance. Sleep 39, 1691–1700, doi:10.5665/sleep.6094 (2016).

31 Chung, S. et al. Identification of preoptic sleep neurons using retrograde labelling and gene profiling. Nature 545, 477–481, doi:10.1038/nature22350 (2017).

32 Lambiase, M. J., Gabriel, K. P., Kuller, L. H. & Matthews, K. A. Temporal relationships between physical activity and sleep in older women. Med Sci Sports Exerc 45, 2362–2368, doi:10.1249/MSS.0b013e31829e4cea (2013).

33 Kline, C. E. The bidirectional relationship between exercise and sleep: Implications for exercise adherence and sleep improvement. Am J Lifestyle Med 8, 375–379, doi:10.1177/1559827614544437 (2014).

34 Paul, K. N., Dugovic, C., Turek, F. W. & Laposky, A. D. Diurnal sex differences in the sleep-wake cycle of mice are dependent on gonadal function. Sleep 29, 1211–1223, doi:10.1093/sleep/29.9.1211 (2006).

35 Campos-Beltran, D. & Marshall, L. Changes in sleep EEG with aging in humans and rodents. Pflugers Arch 473, 841–851, doi:10.1007/s00424-021-02545-y (2021).

36 Whittaker, D. E. et al. A recessive PRDM13 mutation results in congenital hypogonadotropic hypogonadism and cerebellar hypoplasia. J Clin Invest 131, doi:10.1172/JCI141587 (2021).

37 Jorgensen, C. S. et al. Identification of genetic loci associated with nocturnal enuresis: a genome-wide association study. Lancet Child Adolesc Health 5, 201–209, doi:10.1016/S2352-4642(20)30350-3 (2021).

38 Watanabe, S. et al. Prdm13 regulates subtype specification of retinal amacrine interneurons and modulates visual sensitivity. J Neurosci 35, 8004–8020, doi:10.1523/JNEUROSCI.0089-15.2015 (2015).

39 Mizuno-Iijima, S. et al. Efficient production of large deletion and gene fragment knock-in mice mediated by genome editing with Cas9-mouse Cdt1 in mouse zygotes. Methods 191, 23–31, doi:10.1016/j.ymeth.2020.04.007 (2021).

40 Kagoshima, H. et al. The C. elegans CBFbeta homologue BRO-1 interacts with the Runx factor, RNT-1, to promote stem cell proliferation and self-renewal. Development 134, 3905–3915, doi:10.1242/dev.008276 (2007).

41 Sato, Y. et al. A mutation in transcription factor MAFB causes Focal Segmental Glomerulosclerosis with Duane Retraction Syndrome. Kidney Int 94, 396–407, doi:10.1016/j.kint.2018.02.025 (2018).

42 Franken, P., Dijk, D. J., Tobler, I. & Borbely, A. A. Sleep deprivation in rats: effects on EEG power spectra, vigilance states, and cortical temperature. Am J Physiol 261, R198–208, doi:10.1152/ajpregu.1991.261.1.R198 (1991).

43 Tada, H. et al. Neonatal isolation augments social dominance by altering actin dynamics in the medial prefrontal cortex. Proc Natl Acad Sci U S A 113, E7097–E7105, doi:10.1073/pnas.1606351113 (2016).

44 Satoh, A. et al. SIRT1 promotes the central adaptive response to diet restriction through activation of the dorsomedial and lateral nuclei of the hypothalamus. J Neurosci 30, 10220–10232, doi:10.1523/JNEUROSCI.1385-10.2010 (2010).

45 Nakamura, H. et al. Cooperative role of the RNA-binding proteins Hzf and HuR in p53 activation. Mol Cell Biol 31, 1997–2009, doi:10.1128/MCB.01424-10 (2011).

