## Supplementary Figures for "Age-associated sleep-wake patterns are altered with Prdm13 signaling in the dorsomedial hypothalamus and dietary restriction in mice"

[1]Department of Integrative Physiology, [4]Department of Inflammation and Immunosenescence, Geroscience Research Center, [8]Laboratory of Experimental Animals, National Center for Geriatrics and Gerontology (NCGG), Obu, Japan.
[2]Department of Developmental Biology, [6]Department of Neurology, Washington University School of Medicine, St. Louis, MO, USA.
[3]Department of Nutrition, Faculty of Wellness, Shigakkan University, Obu, Japan.
[5]Department of Physiology, Yokohama City University Graduate School of Medicine, Yokohama, Japan.
[7]Laboratory Animal Resource Center, University of Tsukuba, Tsukuba, Ibaraki, Japan.
[9]Department of Pathology and Biological Responses, [10]Department of Integrative Physiology, Nagoya University Graduate School of Medicine, Nagoya, Japan.
[11]Department of Gerontology, Laboratory of Molecular Life Science, Institute of Biomedical Research and Innovation (IBRI), Kobe, Japan.
[12]Department of Integrative Physiology, Institute of Development, Aging, and Cancer, Tohoku University, Sendai, Japan.
[13]equally contributing authors

[*]Corresponding author:
Akiko Satoh, Ph.D.
Associate Professor
Department of Integrative Physiology
Geroscience Research Center,
National Center for Geriatrics and Gerontology
7-430 Morioka-cho, Obu, Aichi, 474-8511, Japan


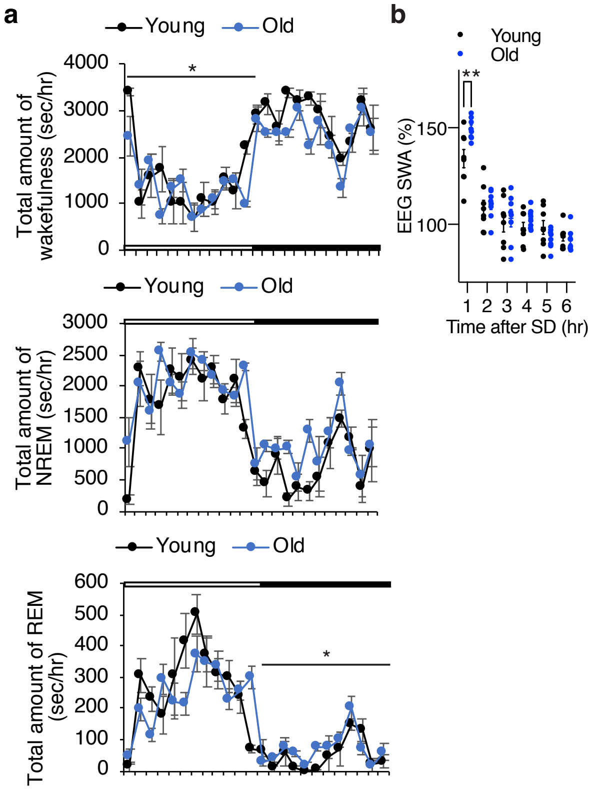


**Supplementary Fig. 1: Old C57BL/6J mice display increased sleep propensity and increased SWS after SD. a**, Total amount of wakefulness (top), NREM sleep (middle) and REM sleep (bottom) in young and old mice. Open bar indicates the light period and filled bar indicates the dark period (n=8). Values are shown as means ± S.E., *p<0.05 by repeated measures ANOVA. **b,** SWA after SD. Each value is relative to the average of the 24-hour baseline day. Values are shown as means ± S.E., **p<0.01, by repeated measures ANOVA with Bonferroni's post hoc test.

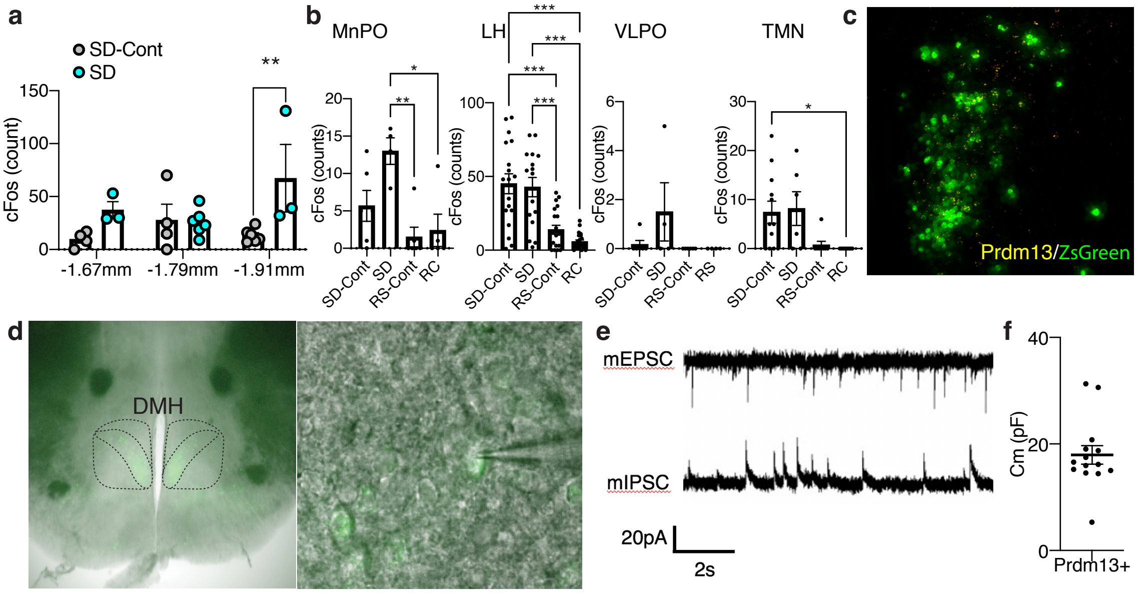


**Supplementary Fig. 2: Neurons are activated in the DMH during SD, and Prdm13+ cells are electrically active. a**, Numbers of cFos+ cells in the DMH at bregma -1.67, -1.79 and -1.91 mm during SD and sleeping-control (SD-Cont) detected by cFos immunohistochemistry (n=3-6). Values are shown as means ± S.E., **p<0.01 by two-way ANOVA with Bonferroni's post hoc test. **b**, Numbers of cFos+ cells in the median preoptic area (MnPO), lateral hypothalamus (LH), ventromedial preoptic area (VLPO) and tuberomammillary nucleus (TMN) during SD, recovery sleep (RS) and sleeping-control (SD-Cont and RS-Cont) detected by cFos immunohistochemistry (n=4-22). Values are shown as means ± S.E., *p<0.05, **p<0.01 and ***p<0.001 by one-way ANOVA with Bonferroni's post hoc test. **c**, Representative image of the DMH with ZsGreen fluorescence derived from *Prdm13*-CreERT2 (green) and mRNA of *Prdm13* (yellow) visualized by RNAscope. **d,** Representative images showing Prdm13+ neurons in the DMH (left). High-magnification images of Prdm13+ DMH cells with rectangles (right). **e,** Representative mEPSC and mIPSC traces obtained from Prdm13+ DMH cells. **f**, Membrane capacitance of Prdm13+ DMH cells recorded in whole-cell patch (n=14). Values are shown as means ± S.E.

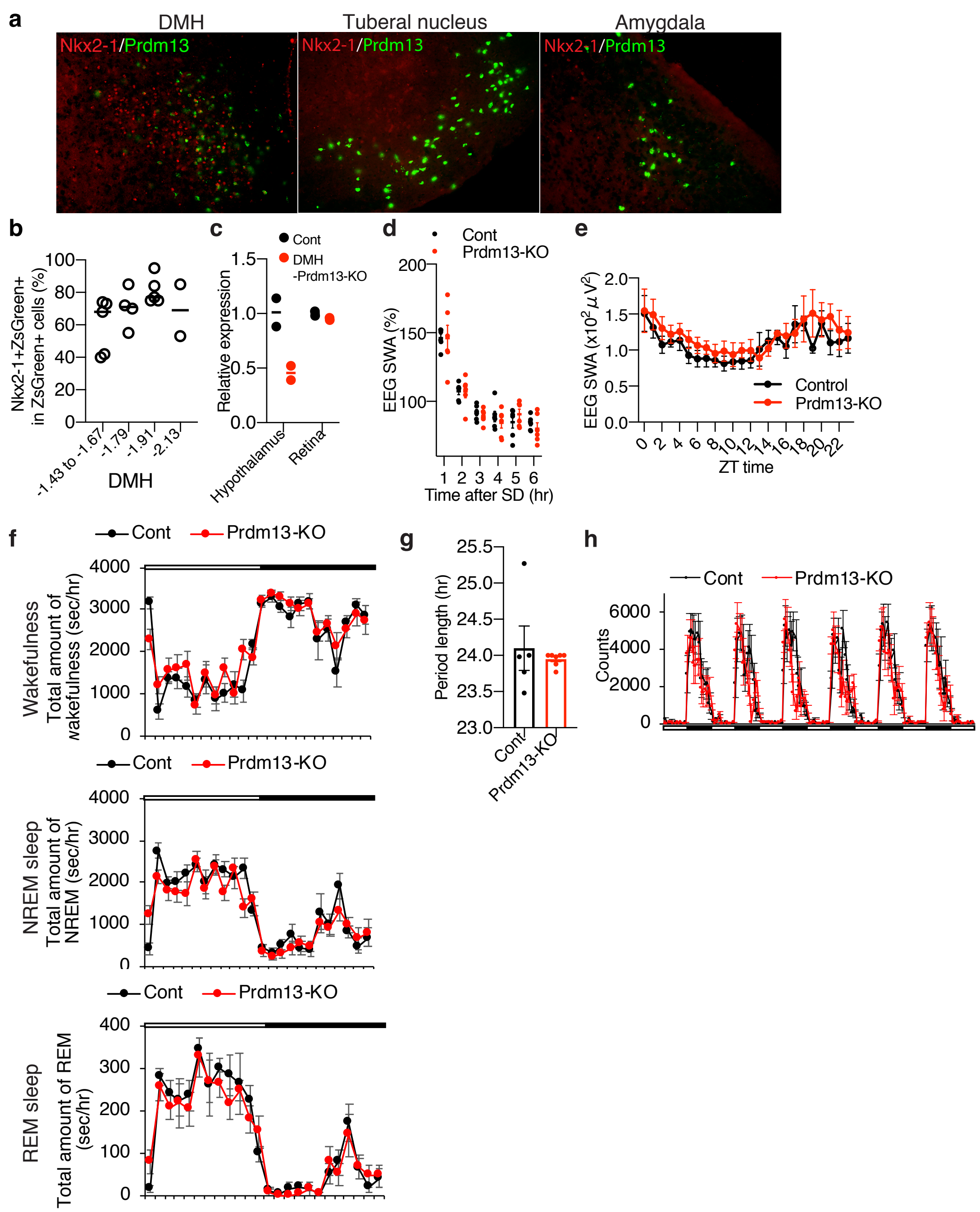


**Supplementary Fig. 3: DMH-specific *Prdm13*-knockout mice do not display changes in sleep/wakefulness architecture, period length and physical activity at young age. a**, Immunohistochemistry of Nkx2-1 (red) with ZsGreen fluorescence derived from *Prdm13*-CreERT2 (green) in the DMH, tuberal nucleus and amygdala. **b**, Percentage of Nkx2-1+ cells within ZsGreen+ cells in the DMH at Bregma -1.43 to -1.67, -1.79, -1.91 and -2.13 mm (n=2-5). Values are shown as means. **c**, Expression of *Prdm13* in the hypothalamus and retina of DMH-specific *Prdm13*-knockout (Prdm13-KO) and control (Cont) mice (n=2). Values are shown as means. **d**, SWA during NREM sleep after SD. Normalized power is relative to the average of the 24-hour baseline day each group (n=6). Values are shown as means ± S.E. **e,** SWA in the range of frequencies between 0.5 to 4 Hz during NREM sleep for a 24-hour period (n=5). Normalized power is relative to the average of the 24-hour baseline day each group. Values are shown as means ± S.E. **f**, Total amount of wakefulness (top), NREM sleep (middle) and REM sleep (bottom) in Prdm13-KO and Cont mice. Open bar indicates light period and filled bar indicates dark period (n=6). Values are shown as means ± S.E. **g**, Period length of Prdm13-KO and Cont mice (n=5-7). Values are shown as means ± S.E. **h**, The level of wheel-running activity in Prdm13-KO and Cont mice for six consecutive days (n=5-7). Values are shown as means ± S.E.

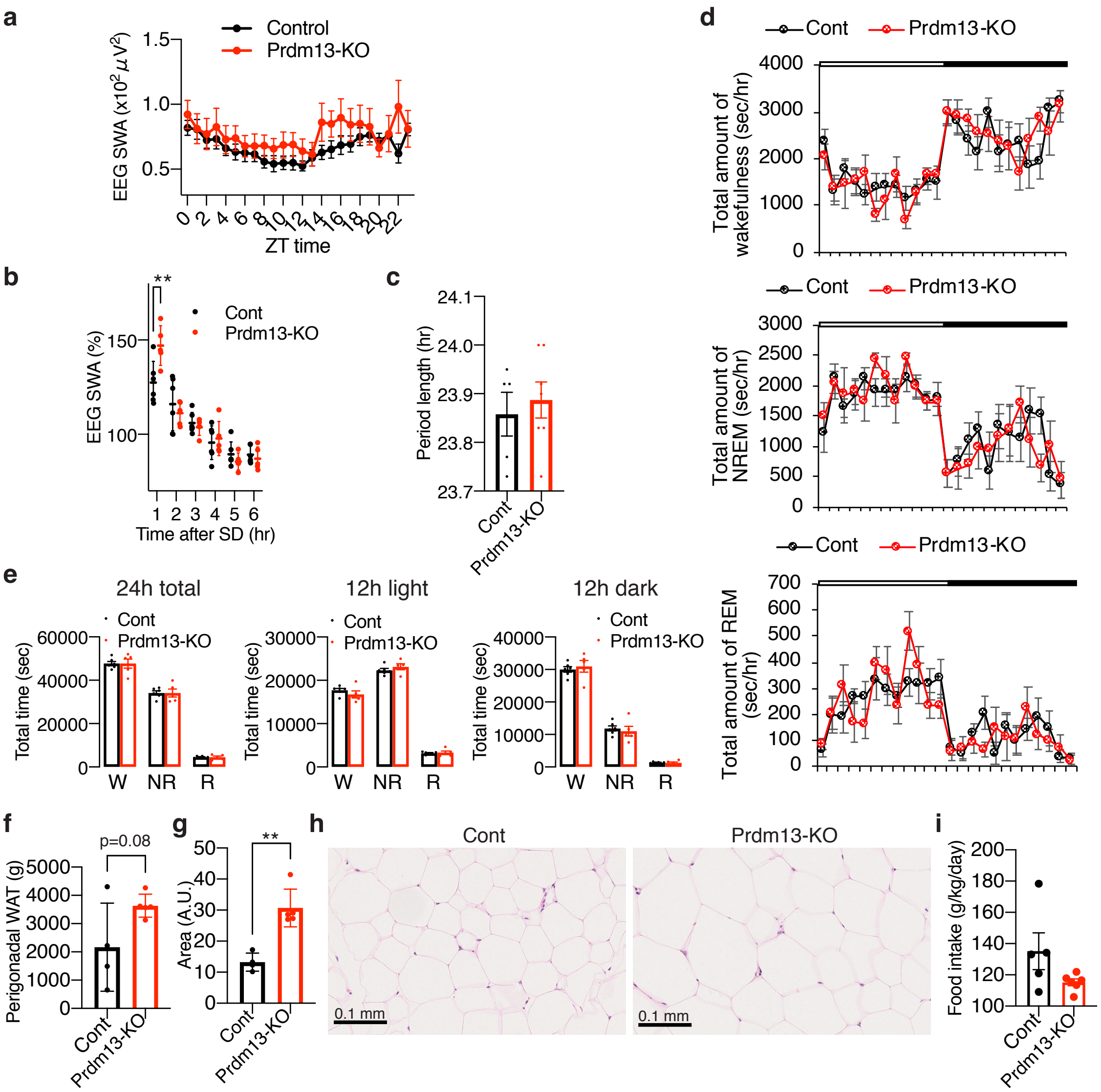


**Supplementary Fig. 4: Old DMH-specific *Prdm13*-knockout mice display high SWA after SD, no abnormality in the amount of sleep/ wakefulness but increased adipose tissue. a,** SWA in the range of frequencies between 0.5 to 4 Hz during NREM sleep for a 24-hour period (n=5-6) in old DMH-specific *Prdm13*-knockout (Prdm13-KO) and control (Cont) mice. Normalized power is relative to the average of the 24-hour baseline day each group. Values are shown as means ± S.E. **b,** SWA after SD. Normalized power is relative to the average of the 24-hour baseline day. Values are shown as means ± S.E. **c**, Period length of old Prdm13-KO and Cont mice (n=5-7). Values are shown as means ± S.E. **d,** Total amount of wakefulness (top), NREM sleep (middle) and REM sleep (bottom) in old Prdm13-KO and Cont mice. Open bar indicates light period and filled bar indicates dark period (n=5-6). Values are shown as means ± S.E. **e**, Total amount of wakefulness (W), NREM sleep (NR) and REM sleep (R) during a 24-hour period (24h total), 12-hour light period (12h light) or 12-hour dark period (12h dark) (n=5-6). Values are shown as means ± S.E. **f**, Weight of the perigonadal WAT in old Prdm13-KO and Cont mice. Values are shown as means ± S.E., listed p-value was calculated by unpaired t-test. **g,h**, Size of adipocytes in the perigonadal WAT in old Prdm13-KO and Cont mice (n=4-5, average of five randomly selected areas). Values are shown as means ± S.E., **$p<0.01$ by unpaired t-test. Representative images are shown in **h**. **i**, Food intake of old Prdm13-KO and Cont mice (n=5-6). Values are shown as means ± S.E.

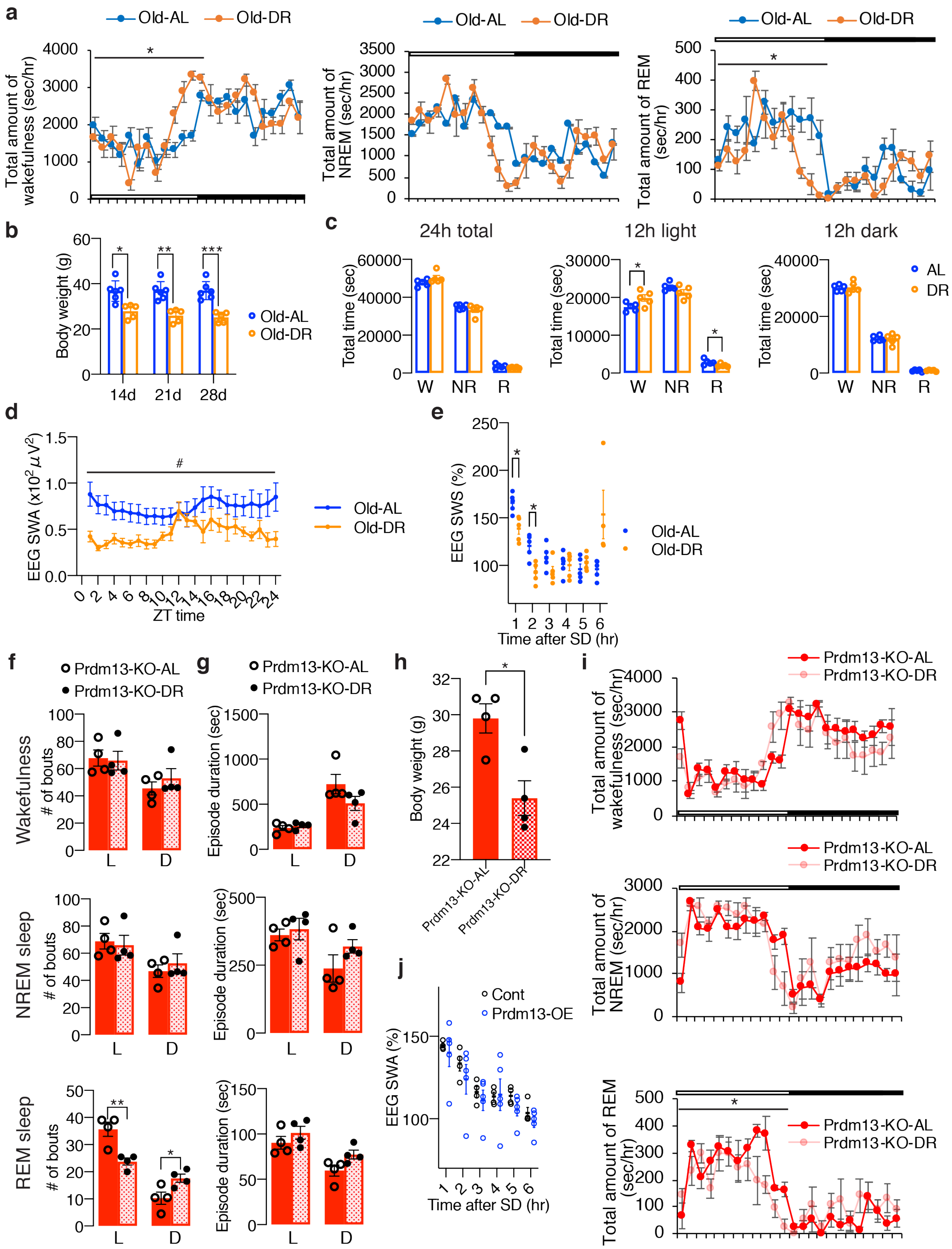


**Supplementary Fig. 5: Diet restriction and overexpression of *Prdm13* in the DMH ameliorates excessive sleepiness during SD in old mice. a**, Total amount of wakefulness (left), NREM sleep (middle) and REM sleep (right) in old-diet restricted (Old-DR) and old-*ad libitum* (Old-AL) mice (n=5). Open bar indicates light period and filled bar indicates dark period. Values are shown as means ± S.E., *p<0.05 by repeated measures ANOVA.

**b**, Body weight of Old-DR and Old-AL mice for 14, 21 and 28 days (n=5-6). Values are shown as means ± S.E., *p<0.05, **p<0.01 and ***p<0.001 by repeated measures AVOVA with Bonferroni's post hoc test. **c,** Total amount of wakefulness, NREM sleep and REM sleep during a 24-hour period (24h total), 12-hour light period (12h light) or 12-hour dark period (12h dark) (n=5). Values are shown as means ± S.E., *p<0.05 by unpaired t-test. **d,** SWA after SD of Old-AL and Old-DR mice. Normalized power is relative to the average of the 24-hour baseline day (n=5). Values are shown as means ± S.E., *p<0.01 by Bonferroni's post hoc test. **e,** SWA during NREM sleep for a 24-hour period. Normalized power is relative to the average of the 24-hour baseline day (n=5-6). Values are shown as means ± S.E., #p<0.05 by repeated measures ANOVA. **f,g**, Numbers of episode (**f**) and duration (**g**) of wakefulness (top), NREM sleep (middle) and REM sleep (bottom) during the light (L) and dark (D) periods in AL and DR of DMH-specific *Prdm13*-knockout (Prdm13-KO-AL and Prdm13-KO-DR, respectively) mice (n=4). Values are shown as means ± S.E., *p<0.05 and **p<0.01 by unpaired t-test. **h,** Body weight of Prdm13-KO-AL and Prdm13-KO-DR mice (n=4). Values are shown as means ± S.E., *p<0.05 by unpaired t-test. **i**, Total amount of wakefulness (top), NREM sleep (middle), and REM sleep (bottom) in Prdm13-KO-AL and Prdm13-KO-DR mice (n=4). Open bar indicates light period and filled bar indicates dark period. Values are shown as means ± S.E., *p<0.05 by repeated measures ANOVA. **j**, SWA after SD of Prdm13-OE and control (Cont) mice. Normalized power is relative to the average of the 24-hour baseline day (n=4). Values are shown as means ± S.E.

**a**

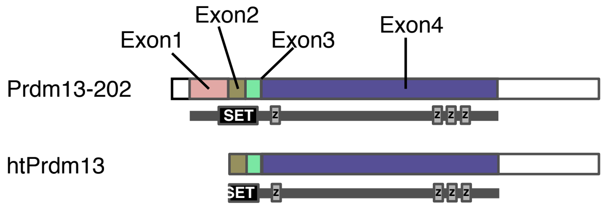


**b**

```
ATGGTGAGAGGGGAGCTGGTGGACGAGTCAGGGGGCTCCCCTCTGGAGTGGATAGGGTTAATCCGGGCAGCCAGGAATCCCCAGGAACAGACTCTGG
AAGCTATTGCAGACTTACCAGGAGGACAGATCTTCTATCGGGCATTGCGAGACGTCCAGCCTGGGGAAGAGCTGACTGTGTGGTATTCTAACTCCTT
GGCTCAGTGGTTTGACATCCCTACAACTGCGACTCCAACGCATGATGAGAAAGGGGAGGAGCGCTACATCTGCTGGTACTGCTGGAGGACCTTTCGC
TACCCTAACAGCCTCAAGGCACACCTGCGGTTCCACTGCGTCCTCAGTGGCGGCGGGGGCCGTGCTTTCCTGCCCCAGGAGCACGCGGCCCGGCCCG
GCGCTTCCCCGGTGGCAGAAGGCCTCGGCCTCCCTCCGAAACCCACCGTCCCGGACTTGACGGCGCCCGTCCAGGCAATCGCTTTGCGACCCCAAGC
TCCGGCCGCTCAGCTCGCCCAGGCGTGCGGGGCGCGGGAGAGCATCAAGCGGGAGGCTTCCCTGGCGCCCCTGGCCACCTCCCCGCCGCCGGGCAAG
TGGGGAACGCCCAAGAAGGGCAAGGAGCAGCCGGACCGCGCGCACAGTCAATTCCTGGGGATAGTGGGCGGCTCCTCGGGCGGCGGCGGAGGCCTGC
CATTCTACCCAGGCGTGCGCTCGGCTTTCAAGCCGGCCGGCCTGGCCCGGGCGGCTGCGCAGAGCGACCCCTACCGGGAAGAGGGCGGCGGCAAGGG
ACCGGGGCTGGCGCTAGGGCGGCTGCTGGGCGGGGGCCGGGCCGGCGGGCGTCCGGGCAGTGGGGAGAGCCCGGCGGGCCACCACCACCATCACCAC
CACGCTCACCACCACCACCACCACCATCCCAAGTGCCTGCTGGCCGGGGAGCCGCCGCCGGCGGGCCTGCCGTGTCCCGGGGCCCTGAGAGCCTTCC
CCCTGCTCGCCGGCCACCCGGAAGAGGCGTCGGCCTTCAAGCACGTGGAGCGGGCTCCGCCCGCCGCCGCCACCACCTCCCTGCCCTCCGCGCGCTA
CGCCGCTCTCCCGGCGCCCGGGCTCCCTGTGGAGCGCTGCGCGCTGCAGCCCCTCGACGGCGGCAGCCTCAAGGCTTACCCCGGCGGCGGTGGCGGC
GGCGAGTGCAGCCCCCTGCCCGCCGTCATGCCGGCCTTCACGGTGTACAGCGGAGACCTGCTCTACGGCCCTCCGGCCGCCTATTACCCTCTCAAGC
TGCACTTGGGCGGGCTGCTCAAATACCCCGAGTCCATCTCCTATCTGAGTGGCCCCGCGGCGGCAGCGGCGGCGGCGGCAGCGGCAGCGGCGGCGGC
GGCGGCGATCGGGCCGGCCGAACTGGGCTCCCTGGCCAGCATCGACCGCGAGATCGCCATGCACACACAGCAGCTGTCCGAGATGGCAGCCGGGAAG
AGTCGCGCCCGCCTGGACTCGGGGACGCTGCCTCCAGCCGTAGTGGCGGCGACGGGCCCCGGGGGTGGCGGCGGTGGCGGGAGCGCGGCAGGCAAGC
CCAAGACAGGCCACTTGTGCCTCTACTGTGGCAAGCTGTACTCGCGCAAGTACGGGCTCAAGATCCACATGCGGACGCACACGGGCTACAAGCCGCT
CAAGTGCAAAGTGTGCCTGCGGCCCTTCGGCGACCCCAGCAATCTCAACAAGCACATCCGACTCCACGCCGAGGGCAATACGCCCTACCGCTGCGAG
TTCTGCGGCAAGGTGCTGGTGCGCCGCCGAGACCTGGAGCGTCACGTCAAGTCCCGCCACCCAGGCCAGAGCCTGATGGCCAAGGCGGGCGACGGCC
CGGGCCCGGAGCCCAGCTATGCTCTGGAGCCTGGGGATCCCAAGAGCGAGGACAGTGATGTGGATGTCTGCTTCACTGACGATCAGAGCGACCCCGA
GGCTGGGGGTCGTGGCGAGCACGACTCCTAA
```

```
MVRGELVDESGGSPLEWIGLIRAARNPQEQTLEAIADLPGGQIFYRALRDVQPGEELTVWYSNSLAQWFDIPTTATPTHDEKGEERYICWYCWRTFR
YPNSLKAHLRFHCVLSGGGGRAFLPQEHAARPGASPVAEGLGLPPKPTVPDLTAPVQAIALRPQAPAAQLAQACGARESIKREASLAPLATSPPPGK
WGTPKKGKEQPDRAHSQFLGIVGGSSGGGGGLPFYPGVRSAFKPAGLARAAAQSDPYREEGGGKGPGLALGRLLGGGRAGGRPGSGESPAGHHHHHH
HAHHHHHHHPKCLLAGEPPPAGLPCPGALRAFPLLAGHPEEASAFKHVERAPPAAATTSLPSARYAALPAPGLPVERCALQPLDGGSLKAYPGGGGG
GECSPLPAVMPAFTVYSGDLLYGPPAAYYPLKLHLGGLLKYPESISYLSGPAAAAAAAAAAAAAAAIGPAELGSLASIDREIAMHTQQLSEMAAGK
SRARLDSGTLPPAVVAATGPGGGGGGGSAAGKPKTGHLCLYCGKLYSRKYGLKIHMRTHTGYKPLKCKVCLRPFGDPSNLNKHIRLHAEGNTPYRCE
FCGKVLVRRRDLERHVKSRHPGQSLMAKAGDGPGPEPSYALEPGDPKSEDSDVDVCFTDDQSDPEAGGRGEHDS*
```

**Supplementary Fig. 6: Structures of Prdm13-202 and hypothalamic Prdm13. a**, Prdm13-202 is composed of a SET-domain (SET) and four zinc-finger domains (z) within four exons (from Exon 1 to Exon 4). A proximal portion of SET is deleted in hypothalamic Prdm13 (htPrdm13). **b**, The sequences of base (top) and amino acid (bottom) for htPrdm13.

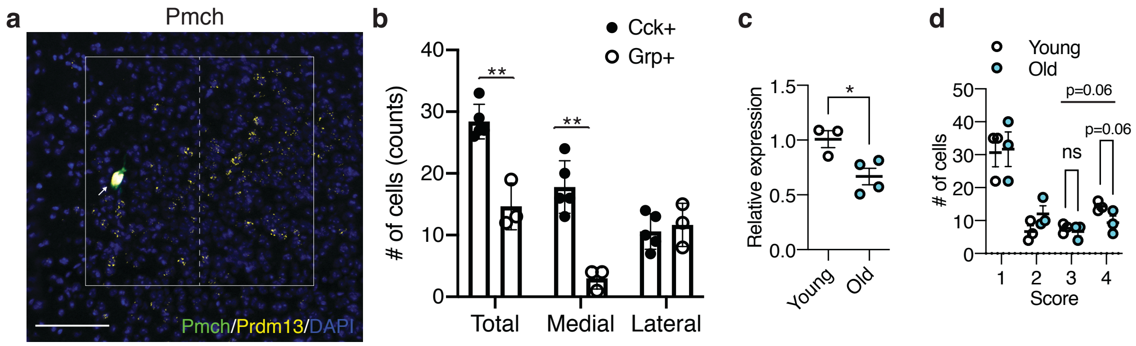


**Supplementary Fig. 7: Distribution of *Pmch* in the DMH, and expression of *Cck* in young and old mice. a**, Representative image of the DMH with *Prdm13* (yellow) and *Pmch* (green) visualized by RNAscope. Cells were counterstained with DAPI (blue). White box shows the DMH, which is divided into medial and lateral areas by dashed line. White arrow shows yellow+ and green+ cells. Scale bar indicates 100 $\mu$m. **b**, Number of *Cck*+ or *Grp*+ cells within *Prdm13*+ cells in medial, lateral or total DMH (n=3-5). Values are shown as means ± S.E., **p<0.01 by unpaired t-test. **c,** Expression of *Cck* mRNA in the hypothalamus of young and old mice (n=3-4). Values are shown as means ± S.E., *p<0.05 by unpaired t-test. **d**, Semiquantitative evaluation of *Cck* expression levels in *Prdm13*+*Cck*+ cells in young or old mice during SD-Cont. Signal dots derived from *Cck* mRNA were counted manually and categorized into four grades: 1 (1-5 dots/cell), 2 (6-10 dots/cell), 3 (11-15 dots/cell) and 4 (>16 dots/cell). Numbers of cells divided into four grades of *Cck* expression were shown as means ± S.E., listed p-values and non-significant (ns) by two-way ANOVA with Bonferroni's post hoc test.
